# A parsimonious model for mass-univariate vertex-wise analysis

**DOI:** 10.1101/2021.01.22.427735

**Authors:** Baptiste Couvy-Duchesne, Futao Zhang, Kathryn E. Kemper, Julia Sidorenko, Naomi R. Wray, Peter M. Visscher, Olivier Colliot, Jian Yang

## Abstract

Covariance between grey-matter measurements can reflect structural or functional brain networks though it has also been shown to be influenced by confounding factors (e.g. age, head size, scanner), which could lead to lower mapping precision (increased size of associated clusters) and create distal false positives associations in mass-univariate vertex-wise analyses. We evaluated this concern by performing state-of-the-art mass-univariate analyses (general linear model, GLM) on traits simulated from real vertex-wise grey matter data (including cortical and subcortical thickness and surface area). We contrasted the results with those from linear mixed models (LMMs), which have been shown to overcome similar issues in omics association studies. We showed that when performed on a large sample (N=8,662, UK Biobank), GLMs yielded large spatial clusters of significant vertices and greatly inflated false positive rate (Family Wise Error Rate: FWER=1, cluster false discovery rate: FDR>0.6). We showed that LMMs resulted in more parsimonious results: smaller clusters and reduced false positive rate (yet FWER>5% after Bonferroni correction) but at a cost of increased computation. In practice, the parsimony of LMMs results from controlling for the joint effect of all vertices, which prevents local and distal redundant associations from reaching significance. Next, we performed mass-univariate association analyses on five real UKB traits (age, sex, BMI, fluid intelligence and smoking status) and LMM yielded fewer and more localised associations. We identified 19 significant clusters displaying small associations with age, sex and BMI, which suggest a complex architecture of at least dozens of associated areas with those phenotypes.

## 4. Introduction

Brain MRI scans can generate hundreds of thousands of vertex/voxel-wise measurements per individual, which can be linked to other measured traits/diseases using mass univariate vertex/voxel-wise association analyses. Results of association analyses (and subsequent follow-up analyses) can shed light on the brain networks or cell composition relevant for the trait/disease and may be leveraged for brain-feature based phenotype prediction. However, brain measurements may exhibit a pattern of correlation, owing to factors (e.g. head size, MRI scanner/artefact (Chen et al., 2020) or demographics (Montembeault et al., 2012)) which can generate confounded brain-trait associations. Induced local correlations with a true brain-biomarker can generate a smear of association (i.e. a cluster of associated vertices) which may limit the precise localisation of the directly associated regions. On the other hand, long-range vertex correlations caused or inflated by factors irrelevant to the trait of interest, may be more prejudicial, as they can yield distal false positives (**Figure 1**).

**Figure 1:**
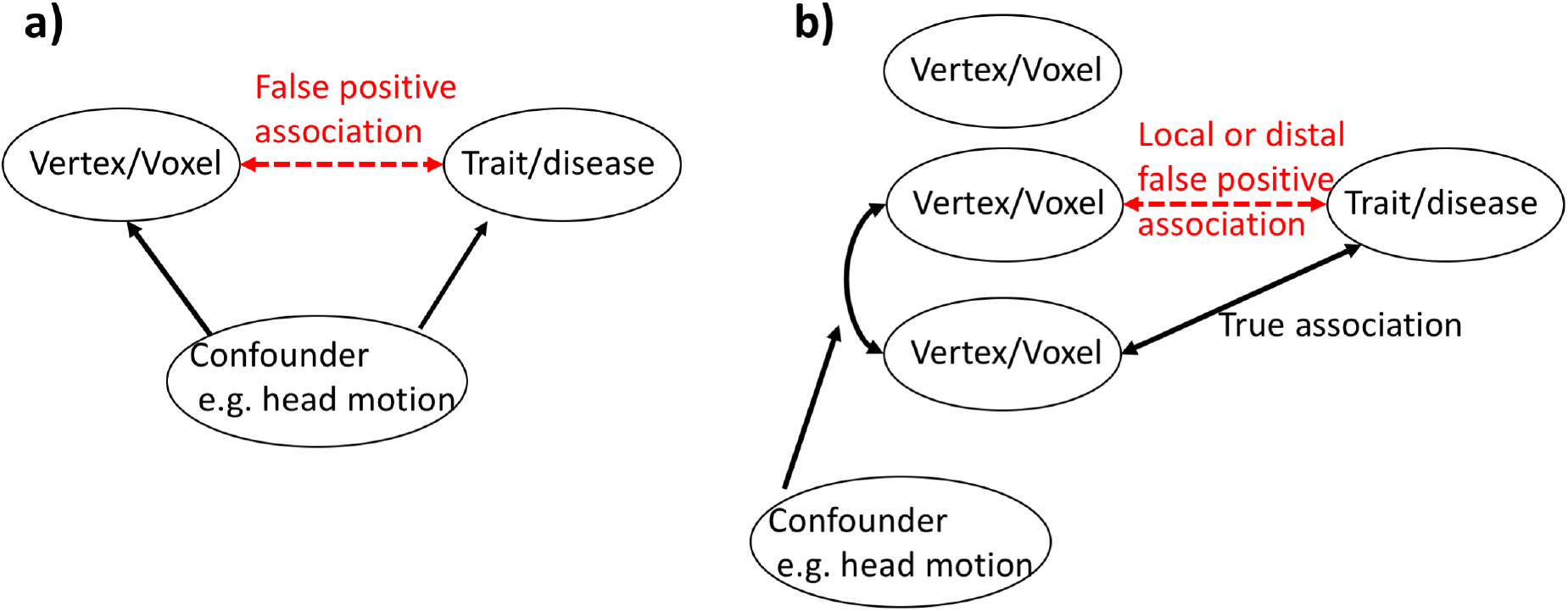
Illustration of the traditional confounding paradigm a) and of the confounding that may arise in association studies performed across correlated brain features b). One sided arrows represent a causal effect, while two-sided arrows represent a correlation.

Two approaches can be used to limit the inflation of false positives described above. One is to control for the confounders in the association testing, although it requires knowledge and measurement of the factors influencing (or more generally associated with) the covariance between brain measurements. Note that these factors can overlap with traditional confounders of neuroimaging studies (e.g. head size, age, sex, head motion), and additional confounders are being identified as sample sizes increase (Alfaro-Almagro et al., 2020). Another correction strategy is to control for the other vertices in the association testing, in order to remove the signal that could be attributed to another brain vertex or region. The difficulty of such approach is that typically, the number of vertex/voxel-wise measurements (p) far exceeds the number of participants (N) in the study. The p>>N paradigm implies that the marginal joint associations with all p vertices cannot be estimated in a single general linear model (GLM).

Statistically, the challenge of mass univariate vertex-wise analyses resembles that of genome-wide association studies (GWAS) or methylation-wide associations studies (MWAS), which aim to identify genomic regions associated with a phenotype in the presence of correlated features (i.e., genetic variants or DNA methylation probes). Several studies have demonstrated that feature correlation (i.e., Linkage Disequilibrium (LD) or population structure in genetics) can result in inflated false positive rate (Cardon & Palmer, 2003; Marchini, Cardon, Phillips, & Donnelly, 2004; Zhang et al., 2019), even more so when the sample size increases (Marchini et al., 2004). This led GLMs to be replaced by linear mixed models (LMMs) (Price, Zaitlen, Reich, & Patterson, 2010; Yang, Zaitlen, Goddard, Visscher, & Price, 2014; Zhang et al., 2019) which co-varies out all features by fitting them as random effects. LMMs have been shown to better control the inflation of false positive associations arising from LD or correlation between probes and to minimise the occurrence of false positives in both GWAS and MWAS (Nabais et al., 2020; Yang et al., 2014; Zhang et al., 2019).

Here, we sought to evaluate whether the inflation of false positives observed with omics dataset is also present in neuroimaging data. In the first part of the analysis, we performed extensive phenotype simulations from real grey-matter data in order to quantify false positive rate as well as statistical power, mapping precision and resulting prediction accuracy achieved from mass-univariate analyses results. We compared the performances of the current state-of-the-art GLMs to that of LMMs inspired by omics association studies. In a second part, we sought to characterise the brain regions associated with real phenotypes (age, sex, BMI, fluid IQ, and smoking status) and to confirm the results obtained on simulated traits. Our analyses relied on thousands of MRI images collected by the UK Biobank (UKB), one of the largest brain imaging initiative (Miller et al., 2016).

## 5. Material and methods

### 5.1. Models of mass-univariate vertex wise analyses

First, we considered five GLMs that differ in term of covariates used when estimating the association (*b_i_*) between the trait and the *i*th (standardised) vertex-wise measurement (***X_i_***). They can be written under the form:

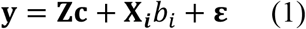

with **y** the vector of phenotype for the **N** individuals, **Z** a matrix of size Nxq of q covariates and ***c*** a vector of the q fixed effects.

The five GLMs are differentiated as follows: 1) GLM with no covariates (“no covariates”), 2) GLM including the most commonly used covariates in similar analyses: age, sex and intra-cranial volume (ICV) (“age, sex, ICV corrected”), 3) & 4) GLMs including 5 and 10 principal components (PCs), respectively (“5 global PCs”, “10 global PCs”), 5) GLM including 10 PCs specific to the measurement type (cortical thickness, cortical surface, subcortical thickness or subcortical surface area), referred to as “10 modality specific PCs”. Grey-matter PCs capture the main axes of covariations between vertices, and we expect that by controlling for them we may be able to remove unmeasured or unknown factors contributing to long-range correlation between vertices (which might include demographics, MRI machine, head motion, software update, processing option *etc.*). Note that PCs from genetic data are commonly used in GWAS in order to limit the false positive rate of GLMs analyses (Price et al., 2006) but are rarely used in neuroimaging analyses. The difficulties of PC correction are to determine the optimal number of PCs and ensuring PCs do not remove signal of interest. In practice, GLMs without covariates are also very rare, but worth considering in order to appreciate the effect of including covariates.

Finally, we considered two LMMs that can be seen as extensions of the previous approaches in that they further control for all vertex-wise measurements. The first LMM model (“LMM global BRM”), analogous to the MOA (MLM-based Omic Association) model (Zhang et al., 2019), can be written as:

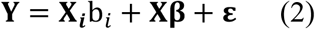

Here, ***X*** is the Nxp matrix of all standardised vertex-wise measurements, **β** is the px1vector of joint vertex-trait associations. **β** is a vector of random effects, allowing for p>N, with 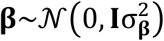, and **ε** is the error term assumed to follow 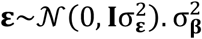 and 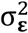 are the variances of the random effects **β** and **ε**. The variance-covariance matrix for **Y** is 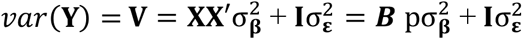. Here, we recognise **B** = **XX**’/p as the brain relatedness matrix and 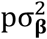 the morphometricity (phenotypic variance captured by the total association with all vertices) (Couvy-Duchesne et al., 2019). Covariates **Zc** may be included in the model but we did not include covariates, in order to evaluate the sole effect of controlling for the random effect (and recognising that covariate variance can be captured by the BRM).

The second LMM (“LMM multi. BRM”) includes 4 random effects (**β_1_ β_2_, β_3_, β_4_**), each corresponding to a type of vertices (cortical thickness, cortical surface area, subcortical thickness and subcortical surface area).

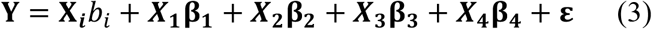

This more general LMM allows the distribution of effect sizes to differ based on vertex type, rather than enforcing a single distribution over all types of measurements (Couvy-Duchesne et al., 2019). Note that each random effect takes up a single degree of freedom meaning that LMMs and GLMs have a comparable (large) numbers of degrees of freedom given the same sample size.

### 5.2. Statistical testing and multiple comparison

We performed a χ^2^ test of the association between a vertex (**X**_*i*_) and the phenotype using that, for large sample size 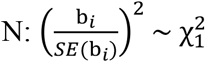 under the null hypothesis of no association. In each model (GLM or LMM), we accounted for multiple testing over the vertices using Bonferroni correction, thus setting a brain-wide significance threshold of 0.05/p=0.05/652,283=7.6e-8. We chose the straightforward Bonferroni correction over random field theory (RFT)(Nichols & Hayasaka, 2003) as RFT requires stationarity and a smooth mesh of vertex-wise residuals which is unlikely to be the case here (we did not apply kernel smoothing on the data as it reduced the estimated morphometricity of the UKB phenotypes (Couvy-Duchesne et al., 2019)). In addition, RFT is not currently implemented to be performed using residuals of LMMs or across several surfaces and type of measurements. Bonferroni correction is expected to be conservative under the null hypothesis (no association) because the correlations between vertices means that the effective number is tests lower than the number of tests conducted and used for the Bonferroni correction.

### 5.3. UKB participants recruitment

The UKB participants were unselected volunteers from the United Kingdom (Sudlow et al., 2015). Exclusion criteria were limited to the presence of metal implant or any recent surgery and health conditions problematic for MRI imaging (e.g. hearing, breathing problems or extreme claustrophobia) (Miller et al., 2016).

### 5.4. T1 weighted and T2 FLAIR image collection

MRI images were mostly collected in Cheadle (for 96% of the sample) and Newcastle using a 3T Siemens Skyra machine (software platform VD13) and a 32-channel head coil (Miller et al., 2016). The T1 weighted (T1w) images were acquired over 4:54 minutes, voxel size 1.0×1.0×1.0mm, matrix of 208×256×256mm, using a 3D MPRAGE sequence, sagittal orientation of slice acquisition, R=2 (in plane acceleration factor), TI/TR=880/2000ms (Miller et al., 2016). The T2 FLAIR acquisition lasted 5:52 minutes, voxel size 1.05×1.0×1.0 mm, matrix of 192×25×256 voxels, 3D SPACE sequence, sagittal orientation, R=2, partial Fourier 7/8, fat saturated, TI/TR=1800/5000ms, elliptical (Miller et al., 2016).

### 5.5. Image processing

We processed the T1w and T2 FLAIR images together to enhance the tissue segmentation in FreeSurfer 6.0 (Fischl, 2012), which should result in a more precise skull stripping and pial surfaces definition. When the T2 FLAIR was not acquired or not usable, we processed the T1w image alone, though a recent report showed this results in systematic differences in cortical thickness (Lindroth et al., 2019). This may represent a source of noise in the data, albeit it was limited in term of number of individuals (see quality control). We extracted vertex-wise data mapping cortical surface area and thickness, using the maximal resolution allowed by the FreeSurfer software (fsaverage - unsmoothed). We previously showed that this cortical processing maximised the morphometricity for a wide range of phenotypes (Couvy-Duchesne et al., 2019). In other words, this cortical processing maximised the information retained by the processed MRI images. In addition, we applied the ENIGMA-shape processing (Boris A. Gutman, Madsen, Toga, & Thompson, 2013; B. A. Gutman, Wang, Rajagopalan, Toga, & Thompson, 2012) to extract radial thickness and log Jacobian determinant (analogous to a surface area (Roshchupkin et al., 2016)) of the hippocampus, putamen, amygdala, thalamus, caudate, pallidum and accumbens. In short, segmented subcortical volume from FreeSurfer are projected onto atlases in order to measure their vertex-wise radial thickness and surface area (Boris A. Gutman et al., 2013; B. A. Gutman et al., 2012). Overall, the imaging data used in the analyses comprised 652,283 vertex measurements per individual: 299,009 for cortical thickness, another 299,034 for cortical surface area, 27,120 for subcortical thickness and 27,120 for subcortical surface area.

In a post-hoc analysis, we also utilised smoothed cortical data (FWHM=20mm), in order to evaluate the robustness of our results to variation in the MRI processing.

### 5.6. Sample description – simulation and discovery sample

We considered the first 10,103 participants of the UK Biobank (UKB) imaging wave. None of the participants withdrew consent after the data were collected. We excluded 231 participants due to T1 images labelled unusable by the UKB or because the FreeSurfer processing failed or did not complete within 48 hours. Our final sample comprised 9,890 adults with complete cortical and subcortical data, aged 62.5 on average (SD=7.5, range 44.6–79.6) with slightly more (52.4%) female participants. To note, 341 participants did not have an exploitable T2 image.

We performed a stringent quality control (QC) of the vertex-wise data in order to exclude participants who may bias the LMM estimates, though there may be a more optimal QC threshold that could maximise the sample size in future studies. Thus, we excluded 1,228 subjects (12.4% of the sample) whose brains were the most similar or dissimilar to that of other individuals (+-5SD from the mean of the brain-relatedness matrix off-diagonal values). Excluding participants with extremely similar brains may seem counterintuitive, though in our experience they tend to be individuals with comparable failed processing (e.g. spike-like cortical parcellation in FreeSurfer)(Couvy-Duchesne et al., 2020). Note that this QC step led to exclude 80.6% of the participants processed using T1w only (vs. 9.9% of the individuals processed using T1w+T2w, p-value<10^-16^), which confirmed our QC could identify individuals or groups of individuals with outlying brain measurements (Lindroth et al., 2019). In addition, QCed out participants had lower cortical thickness and more extreme ICV, cortical thickness, surface area and subcortical volumes (positive associations with the quadratic terms; p<1e-^16^). Finally, our QC excluded slightly more males than females (14.6% vs. 10.4%, p-value=4.6e^-10^) and marginally older participants (63.0 vs. 62.5 years of age, p-value=0.018). Note that we previously showed that in the Human Connectome Project sample, our QC strategy identified a handful of individuals with processing flagged using the ENIGMA visual QC protocol (Couvy-Duchesne et al., 2019).

### 5.7. Independent samples for prediction and replication

Our first independent sample included an additional 4,942 participants of the UKB with a T1w image (downloaded in May 2018, most participants also had an exploitable T2w). The final sample (N=4,160 after processing and QC) was on average 63.1 years old (SD=7.46, range 46.1-80.3) with 52.1% of females.

In addition, we used the OASIS3 (Open Access Series of Imaging Studies) sample (LaMontagne et al., 2019) to evaluate the generalizability of the prediction. The OASIS3 dataset gathers several longitudinal MRI studies conducted in the Washington University Knight Alzheimer Disease Research Center over the past 15 years. Our final sample included 1,006 unique participants after processing based on T1w images and QC. When several visits were available for a participant, we selected the one with the most phenotypic information. Participants were 71.1 years old on average (SD=9.18, range 42.6-95.7) and mostly female (55.5%). Almost a quarter of the participants (23.6%) had a diagnosis of Alzheimer’s disease at the time of imaging.

### 5.8. Mass-univariate analyses on simulated phenotypes

#### 5.8.1. Simulation of phenotypic traits from real grey-matter data

We simulated phenotypic traits from the UKB processed (standardised) grey-matter data. To do so, we randomly selected a set of associated vertices and drew their relative effects from a normal distribution. We then calculated the simulated phenotypes as a linear combination of the individuals’ vertex values and noise (see (Zhang et al., 2019)for formulas). We considered three scenarios that differ in term of number of associated vertices and total association with the phenotype. This global association between grey-matter measurements and a trait has been coined morphometricity (Couvy-Duchesne et al., 2019; Sabuncu et al., 2016) and may be expressed as the proportion of the trait variance (R^2^) captured by the vertex-wise measurement. Our scenarios were: i) 10 associated vertices accounting for a phenotype morphometricity of R^2^=0.20 (i.e. 20% of the trait variance); ii) 100 associated vertices with R^2^=0.50; iii) 1000 vertices with R^2^=40%. For each scenario, we simulated 100 phenotypes.

In follow-up analyses, we simulated phenotypes using the same parameters as above, this time restricting the associated vertices to a single type of measurement. This allowed evaluation of the specificity of each type of measurement, which could possess a unique correlation pattern. In addition, this ensures our phenotypes were not associated with cortical vertices only, which represent 90% of the vertex-wise measurements.

To evaluate the effect of smoothing on our results, we simulated phenotypes from smoothed brain maps (see 5.5). For the ease of computation, we restricted the analysis of smoothed data to the case of 10 associated vertices (R^2^=0.2). We kept the same associated vertices (and weights) as in the previous simulation from unsmoothed data. Finally, we randomly simulated 100 “null” traits, in order to evaluate the calibration of the models under the null hypothesis of no association. All simulations were generated using the OSCA software (Zhang et al., 2019).

#### 5.8.2. Inflation of test statistics

First, we compared the empirical distribution of chi^2^ statistics to the expected distribution, which is assumed to follow a χ^2^(1) for non-associated (null) vertices. We considered the ratio of empirical over expected median chi^2^, known as the inflation factor (λ), which is expected to be equal to one across non-associated vertices. We also used the nominal false positive rate (FPR) defined as the proportion of null vertices with p-values<0.05 (expected to be 0.05). Correlation between associated and null vertices (e.g. due to confounding factors) typically result in an inflation of test statistics, which may cause null vertices to reach significance in mass-univariate analyses.

#### 5.8.3. Power and precision

We quantified the statistical power of the mass-univariate models using the true positive rate (TPR) defined as the proportion or truly associated vertices reaching significance (after Bonferroni correction). In addition, we quantified the mapping precision of mass-univariate analyses by reporting the median size of the true positive (TP) clusters. We defined TP clusters as sets of significant contiguous vertices of the mesh that contain a true positive vertex.

#### 5.8.4. False positives

We reported the Family-Wise Error Rate (FWER) defined as the proportion of replicates with at least one false positive vertex (null vertex significant after Bonferroni correction). In the presence of strong correlation between neighbouring vertices, it is statistically difficult to separate a true positive vertex from the flanking ones, thus we can expect a FWER greater than 5%. Thus, we also reported the cluster FWER defined as the proportion of replicates with at least one false positive cluster. Finally, we reported the proportion of false positive clusters out of all significant clusters (cluster FDR). We labelled false positive clusters, the groups of significantly associated, contiguous vertices that did not contain a true positive association.

In follow up analyses, we simulated associations on a single type of vertex-wise measurements, in order to evaluate the probability of false positive (FWER) arising on the same type of measurements, other types of measurements as well as contra-lateral regions.

#### 5.8.5. Prediction from significant vertices

We evaluated the prediction accuracy achieved from the brain regions reaching significance, in the different mass-univariate models. We used prediction as a meta-criterion to compare the model performances, as it is dependent on power, true and false positives, and association effect sizes. We selected the most significant vertex in each cluster and constructed a linear predictor using association weights (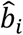, see (1) and (2)) estimated from the different mass-univariate analyses. Because some significant clusters might contain several independent signals, we also built predictors that included all significant vertices. We evaluated the prediction of in the independent UKB and OASIS3 samples.

### 5.9. Mass-univariate analyses of UK Biobank phenotypes

Next, we performed mass-univariate vertex-wise analyses on five UKB phenotypes that showed significant replicated morphometricity (Couvy-Duchesne et al., 2019): age, sex, BMI, smoking status and fluid intelligence. We used the raw fluid intelligence score provided by the UKB, a non-standard test which has demonstrated some reliability in a test-retest analysis (Fawns-Ritchie & Deary, 2020).

For each UKB phenotype and model, we reported the number of significant vertices, number of significant clusters as well as their sizes. We defined significance using a Bonferroni significance threshold of 0.05/(652283*5)=1.5e-8, which accounts for the total number of tests performed. For those phenotypes, the true pattern of association is unknown which prevents evaluation of the false positive rate (or power) of the different approaches. However, false positives or redundant associations should not improve prediction accuracy, which we therefore evaluated for each GLM or LMM model in the UKB replication sample, as well as in the OASIS3 dataset. We used linear predictors as in **section 2.8.5**.

## 6. Results

### 6.1. Analyses on phenotypes simulated under H0 (not associated)

We found that all GLM and LMM models behaved well under the null hypothesis, as indicated by no inflation of test statistic, FPR, or of false positive rate (FWER). As expected under a stringent Bonferroni correction, all approaches were conservative as indicated by FWER<3% (**SFigure 1**).

### 6.2. Analyses on phenotypes simulated under H1 (associated)

#### 6.2.1. Inflation of test statistics

First, we quantified whether we could observe an inflation of test statistics on the vertices not associated with the simulated phenotypes. As expected in presence of correlation between truly associated and null vertices, we observed a global inflation of (median) test statistics when using GLMs (**Figure 2, STable 1**). This was confirmed by an FPR greater than 5% for all GLM models (**STable 1**) even though controlling for covariates or PCs, reduced the inflation of test-statistics compared to the “no covariates” GLM. In comparison, LMMs appropriately controlled the inflation of test statistics on null vertices (λ<1 and FDR<5%; **Figure 2, STable 1**).

**Figure 2:**
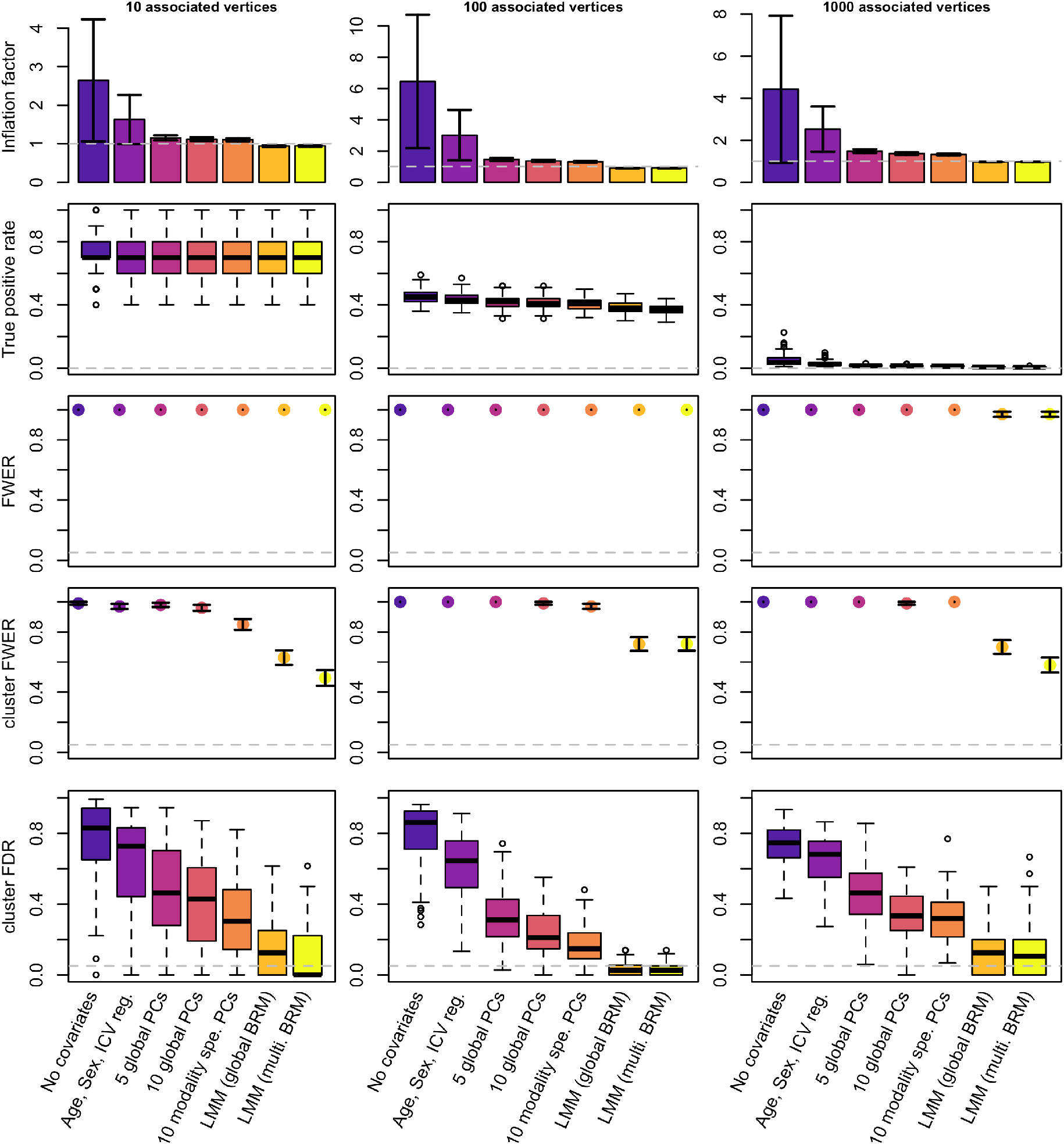
Performance of GLMs and LMMs for mass-univariates vertex-wise analyses: test inflation, statistical power and false positive rate. The columns correspond to the different scenarios considered when simulating traits. We simulated 100 phenotypic traits for each scenario. Bars represent +-SE across the 100 replicates. Clusters are composed of groups of contiguous vertices each significantly associated with the phenotype (after Bonferroni correction). We labelled them false positive if they did not include a true positive association.

#### 6.2.2. Statistical power

First, we confirmed that the statistical power was dependent on the scenarios which corresponded to different effect sizes for the vertices. For example, we had about 70% power to detect associated vertices in the case of a simple trait (10 associated vertices each accounting for 2% of the phenotypic variance on average). On the other hand, statistical power was lower than 5% for the most complex phenotypes (e.g. scenario 3, 1000 vertices each accounting for 0.04% of the variance) (**Figure 2, STable 1**).

Across all scenarios, LMMs exhibited a slightly reduced power compared to the GLMs (**Figure 2, STable1**). We investigated this result using phenotypes simulated from a single type of measurement. We found power of LMMs to be especially reduced on subcortical thickness and surface area (**SFigure 2),** and especially when using the LMM with multiple random effects, which may better control for confounders specific to each vertex type.

#### 6.2.3. False positives

We found that every single simulation yielded at least 1 false positive vertex after Bonferroni correction (FWER=1, **Figure 2**). We noted that the FWER of 0.97 (SE=0.02) found for LMMs in the scenario of “1000 associated vertices”, came from three simulations returning no significant associations. When evaluating the results at a cluster level, we found that using GLMs almost always resulted in one or more false positive cluster (**Figure 2, STable 1**), leading to cluster FWER>85%. Cluster FWER was reduced to 49-72% by using LMMs (**Figure 2, STable 1**). Despite this improvement, no model ensured a cluster-FWER below 5%. LMMs also minimised the proportion of false positive clusters (cluster FDR), compared to the GLM approaches. At the extreme, more than 70% of the significant clusters were false positives using GLMs without covariate. This reduced to about 60% when controlling for age, sex and ICV and further reduced to less than 17% using LMMs (**Figure 2, STable1)**.

Next, we simulated phenotypes associated with a single type of measurement and reported the FWER for each type of measurement in **SFigure 3-6**. This allowed evaluation of whether false positives could appear as a result of associations with vertices from other types of measurements. We found that using GLMs resulted in contamination of signal between all the different types of measurements, as indicated by FWER>5% (**SFigure 3-6**). In comparison, LMMs always minimised the probability of false positives appearing on nonassociated types of measurement. In particular, LMMs ensured that associations on the cortex did not inflate the false positive rate on subcortical structures, and vice versa (FWER<5%, **SFigure 3-6**).

#### 6.2.4. Mapping precision

Here, mapping precision is defined as the size of true positive clusters. LMMs led to a more precise localisation of the associations by minimising the size of true positive clusters (whether we looked a clusters median or maximal size, **Figure 3, STable1**). The median size of true positive clusters was reduced by a factor greater than ten on subcortical measurements, and by a factor greater than two on cortical thickness when using LMMs (**STable 1**). To note, positive clusters on cortical surface area were particularly small (median cluster size of one vertex), independent of the mass-univariate model used, **Figure 3, STable 1**). However, LMMs still offered a greater precision than the GLMs when considering the maximal cluster size (**STable 1**).

**Figure 3:**
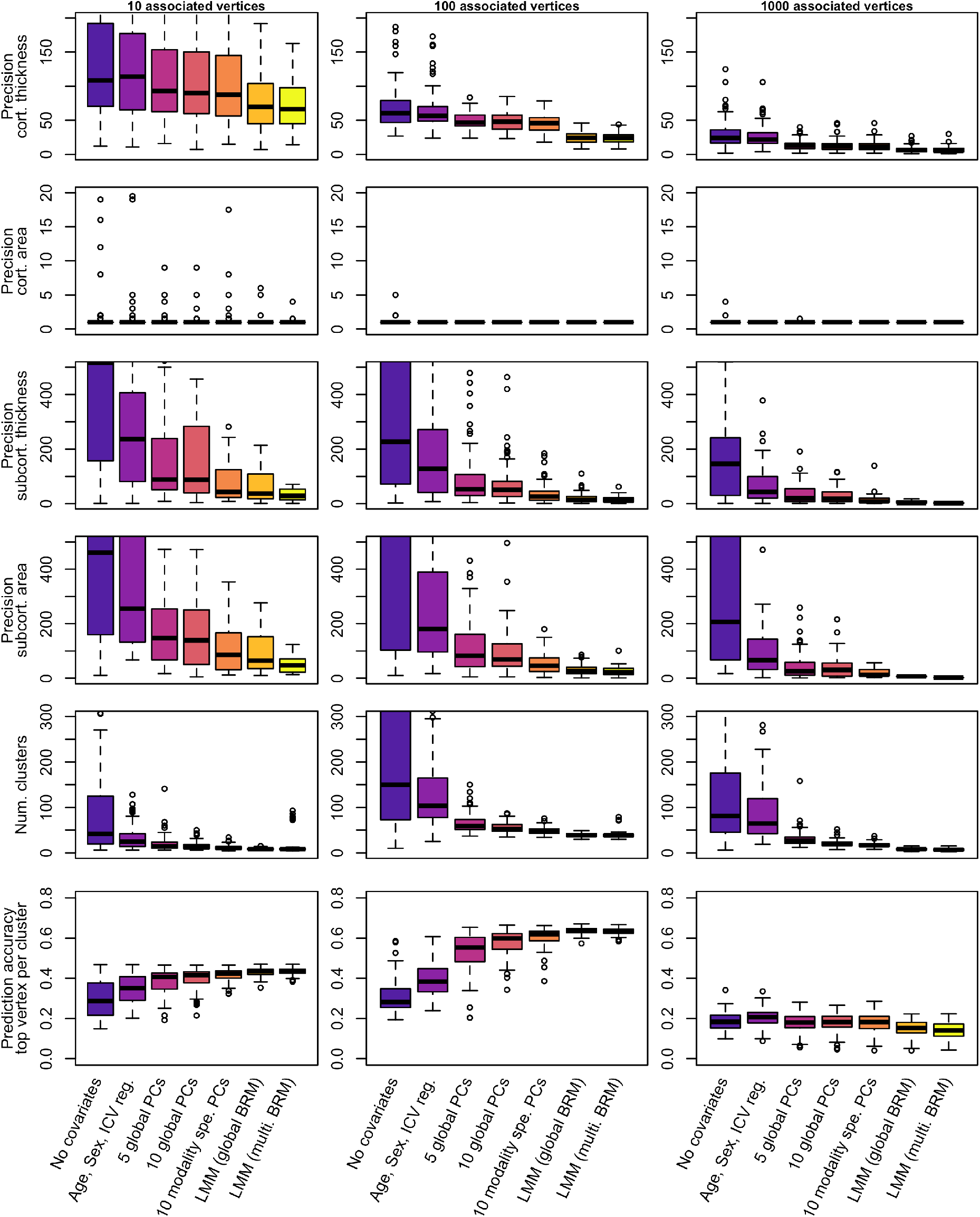
Precision and prediction accuracy from significant vertices between the different models of mass-univariate analyses. The columns correspond to the different scenarios considered when simulating the traits. We simulated 100 phenotypic traits for each scenario. Bars represent +-SE across the 100 replicates. Clusters are composed of groups of contiguous vertices each significantly associated with the phenotype (after Bonferroni correction). We labelled them true positive if they included a true positive association. Precision refers to the median size of true positive clusters.

#### 6.2.5. Prediction accuracy from significant vertices

As a way of aggregating the previous metrics of performance, we compared prediction accuracy achieved from significant vertices, using the UKB replication sample. Across all models and scenarios, selecting the top vertex per significant cluster maximised prediction accuracy, compared to including all significant vertices. This was expected, as significant vertices from the same cluster tag likely redundant information, leading to overweight the prediction signal coming from large clusters.

In simulation scenarios 1 and 2, we found that including more covariates in the GLMs resulted in greater prediction accuracy despite that predictors included fewer vertices (**Figure 3, STable 1**). In addition, LMMs yielded marginally better prediction accuracy than the best GLM using even fewer vertices (**Figure 3, STable 1**), consistent with observation from previous studies (Nabais et al., 2020; Zhang et al., 2019). For the third simulation scenario, the prediction accuracy was comparable and limited for all models (**Figure 3, STable 1**).

#### 6.2.6. Analyses using smoothed cortical surfaces

We repeated the analysis using smoothed cortical meshes of surface and thickness (FWHM=20mm), which is more commonly used in the literature than unsmoothed meshes. We sought to investigate how robust our results were to such variation of MRI processing.

Overall, smoothing did not change the results of the model comparison. LMMs resulted again in a reduced false positive rate (lower cluster FWER and cluster FDR) as well as reduced power (seemingly more important than in the unsmoothed case). LMMs maximised mapping precision and prediction accuracy, despite relying on fewer significant clusters (**SFigure 7**). Of note, performing analyses on smoothed data decreased the mapping precision for the cortex, leading to true positive clusters roughly ten times larger on cortical meshes (**Figure 2, SFigure 7**).

Data smoothing resulted in a large inflation of test statistic and FPR (for GLMs, **Figure 2, SFigure 7**), which is to be expected as smoothing increases the amount of correlation between vertices. We noticed that smoothing led to an increase of cluster FWER for the GLM with 10 PCs, while it decreased cluster FWER for the LMMs (despite the associated vertices and effect sizes remaining the same). This result warrants a more fined-grained evaluation of the associations. We can only hypothesise that the 20mm (FWHM) smoothing can induce medium-range correlations (hence medium range false positives in GLMs) while it also increases local correlation which may aggregate false positive clusters in LMMs.

### 6.3. Morphometricity of the UK Biobank phenotypes

As a proof of concept, we confirmed that the morphometricity of our simulated traits was consistent with that defined in our simulations, whether we fitted a single random-effect component or one random-effect component per modality (**SFigure 8**). For the five UKB phenotypes, we also found consistent morphometricity using both LMM models (**Table 1**), suggesting associations across all types of vertex measurements.

BMI and fluid intelligence exhibited large and moderate morphometricity (R^2^=0.51 (SE=0.031) and R^2^=0.17 (SE=0.034)) but only a limited association with age, sex or the first 10 principal components from vertex-wise data (adjusted R^2^ with ten PCs: R^2^=0.032 for fluid intelligence, R^2^=0.033 for BMI), which resembles the case of our simulations. Age and sex displayed high morphometricity (R^2^=0.83 (SE=0.026) and R^2^=1 (SE=0.024)) and large associations with the first ten PCs (adjusted R^2^=0.41 for age, R^2^=0.43 for sex). Smoking status is a discrete variable (non-smoker, former smoker, still smoking) with a morphometricity of R^2^=0.12 (SE=0.029), and adjusted R^2^=9.2e-3 with first 10 PCs. Note that the morphometricity estimates may be slightly larger than the ones reported previously (Couvy-Duchesne et al., 2019) where we had regressed out age, sex and head size.

### 6.4. Analysis of UK Biobank phenotypes

Using GLM without covariates resulted in many vertices and clusters reaching significance (**Figure 4**, **STable 2**). Unsurprisingly, correcting for covariates which account for a large fraction of the phenotypic variance (see adjusted R^2^ with covariates and PCs, **STable 2**), drastically reduced the number of associations in the GLMs. For example, correcting for ten PCs in mass-univariate analyses of age and sex reduced the number of associated vertices by a factor 8-13, compared to the GLM without covariates (**Figure 4**, **STable 2**). For smoking status, the number of significant vertices and clusters also dropped despite a negligible association with PCs (**Figure 4**, **STable 2**). Similarly, for fluid intelligence, correcting for the top 10 PCs did not remove much of the trait variance over controlling for age sex and ICV (adjusted R^2^=0.030 with age, sex, ICV, adjusted R^2^=0.034 when further controlling for PCs) though it greatly reduced the number of associations, likely because fitting PCs, to some extent, controls for correlations between vertices.

**Figure 4:**
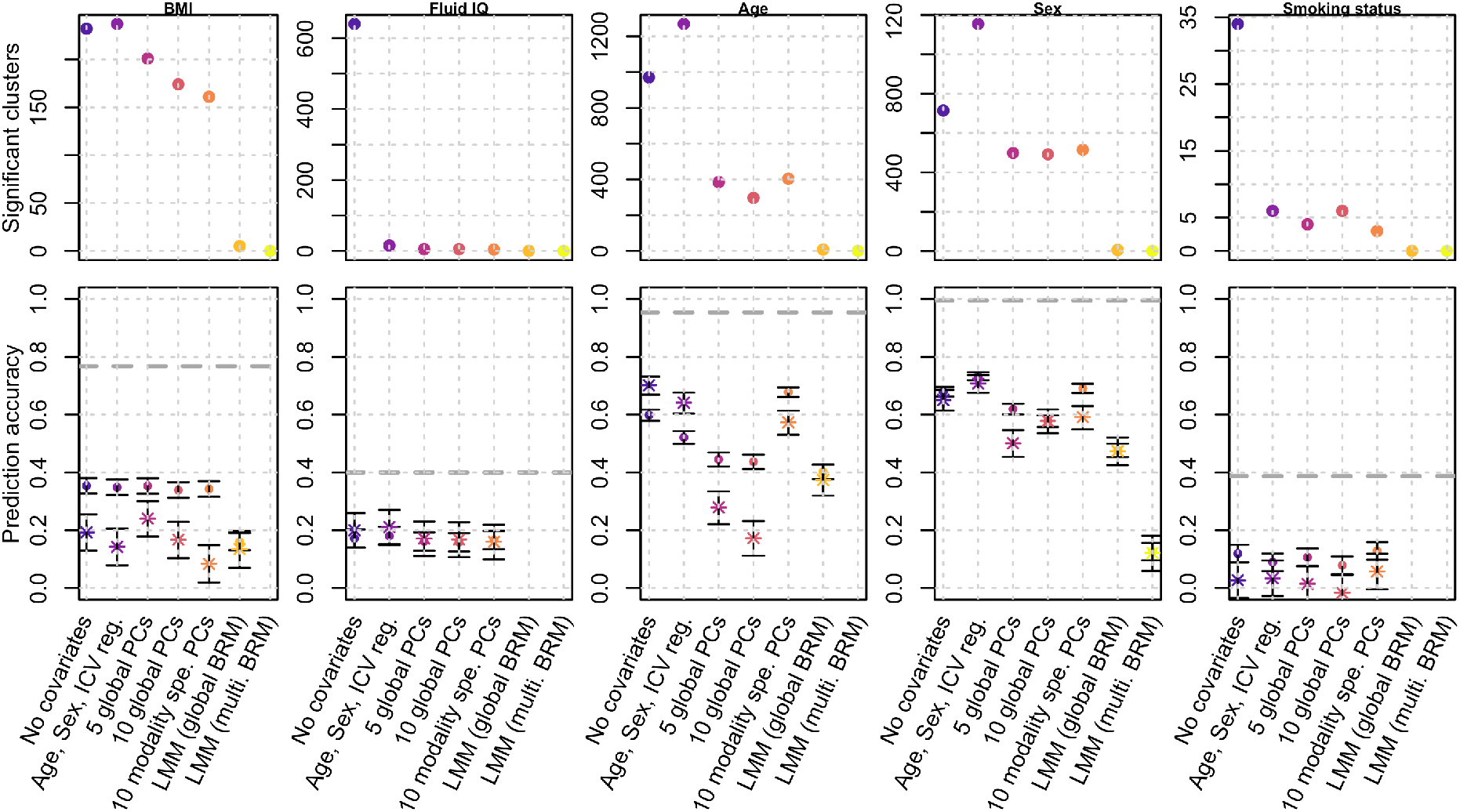
Number of significant clusters and prediction accuracy for real UKB phenotypes. Bars represent the 95% confidence intervals of the prediction accuracy (correlations). Dots indicate prediction accuracy in the UKB replication sample, while stars correspond to the prediction achieved in the OASIS3 sample. The dashed lines correspond to the estimated morphometricity, which corresponds to the theoretical maximum prediction accuracy achievable from a linear predictor.

We found that across all phenotypes, LMMs resulted in a more parsimonious pattern of associations (**Figure 4**, **STable 2**). Thus, with a single random effect LMM, we identified 5 clusters associated with BMI, 8 with age and 6 with sex (**STable 2**). Using an LMM with multiple random-effect components resulted in a single significant cluster for sex reaching significance. Finally, the more covariates (incl. PCs) we corrected for, the smaller the size of the associated clusters, suggesting they do remove confounding effects. LMMs resulted in even sparser associations, consistent with the increased precision observed in simulations. using LMMs (**Figure 4**, **STable 2**).

Next, we compared the prediction accuracy achieved from the vertices reaching significance using each model (**Figure 4**, **STable 2**). Predicting our traits of interest allows evaluating of how power, false and true positive of the different models may counterbalance each other. In addition, prediction into independent samples may serve as an external validation of the findings obtained in the different mass-univariate approaches. For BMI, we found that prediction accuracy from GLMs was greater in the UKB replication sample than in the OASIS3 sample, which suggests that GLMs based predictors capture information that is sample specific (e.g., the same confounders are more likely to be shared in the sample cohort than across different cohorts). In contrast, the prediction accuracy from LMM was comparable on the UKB and OASIS3 samples, pointing towards a better generalisability of such predictor. This also indicates that the higher prediction accuracy in the UKB replication sample for GLM is likely to be driven by the same confounding factor shared between UKB data sets. The comparable performance of GLM and LMM seen on OASIS3 for BMI aligns with our simulations.

For age and sex prediction, prediction accuracy of LMMs was sometimes inferior to that achieved from GLMs, in particular those from the simplest models (“no Covariates” and “age, sex, ICV”). However, prediction from LMM generalised well (comparable accuracy in in UKB and OASIS3), while the GLMs often displayed heterogeneous performances across the test samples (in particular for the GLMs with PCs, which may suffer from PCs being different between samples).

Regarding fluid IQ and smoking status, no LMM predictor was available, and the different GLMs resulted in comparable prediction accuracy, albeit limited in both the UKB and OASIS3 sample.

### 6.5. Description of associated regions from LMM

We summarised the significant associations found using LMM (single modality) in **STable 3 (SFigure9-11** for Manhattan plots, **SFigure 12-16** for brain plots). The significant associations were in the range of R^2^=0.5-1%. Most associations were observed with subcortical volumes though the top cluster for sex was spatially located at the border of the lateral-orbitofrontal and medial orbitofrontal gyri (based on the Desikan atlas(Desikan et al., 2006)). Out of the 85 vertices associated with age, sex and BMI, 68 replicated in an independent UKB sample (p<0.05/85, **Table 2**). In particular, 4/11 associations replicated for BMI, 43/47 with age and with 21/27 sex.

## 7. Discussion

Using simulations, we evaluated the statistical power, false positive rate and precision of GLMs and LMMs for vertex-wise grey-matter association studies. In particular, we evaluated the different models in the context of big-data neuroimaging (large sample size but even greater number of correlated brain vertices) (Smith & Nichols, 2018). We consistently found that using GLMs resulted in a large number of false positive associations and clusters, whether we used smoothed or not-smoothed grey-matter surfaces. Thus, across all scenarios tested, more than 60% of the significant clusters were false positives using a standard GLM that controlled for age, sex and ICV. In comparison, false discovery rate was below 17% using LMMs, though still greater than the 5% expectation (**STable 1, Figure 2, SFigure 7**). In addition, we showed that unlike GLMs, LMMs could appropriately separate cortical from subcortical associations, even though signal contamination between thickness and surface still occurred (**SFigure 2-5).**

Our results suggest that previously reported results from mass univariate vertex-wise analyses obtained using standard GLM approaches could contain many redundant associations, some of which **are likely** to be false positives induced by **confounding factors that cause correlation** between vertices (e.g. (Cox et al., 2019; Navas-Sanchez et al., 2016; Ritchie et al., 2018; Tamnes et al., 2017), see also Figure 1b). Note that albeit redundant in term of association and prediction, some of the brain regions identified using GLM may correspond to indirect manifestations of the trait/disease of interest, which may be relevant to understand the dynamics of grey-matter structure. Importantly, the type 1 error (greater than 5%) we observed in simulations also warns against taking for granted results from LMMs. The increased false positive rate for GLMs has been well documented in omics association analyses studies (e.g. GWAS (Price et al., 2006; Price et al., 2010) or MWAS (Nabais et al., 2020; Zhang et al., 2019)) and has been attributed to proximal and distal correlations between features, caused by factors independent of the trait of interest (e.g. genetic ancestry in genetics, (Price et al., 2006), cell composition of the biological sample and smoking status in DNA methylation (Jaffe & Irizarry, 2014; Zhang et al., 2019)). On the other hand, LMMs can reduce the probability of generating false positives, by fitting all other vertices as random effects which accounts for the complex correlation structure between vertices within and between individuals. In brain imaging, more work is needed to identify the factors that contribute to local and distal correlations between vertices, hence inducing a correlation between true associations and “null” vertices, beyond the usual covariates or confounders used in neuroimaging (e.g. MRI scanner/artefact (Chen et al., 2020) or demographics (Montembeault et al., 2012)).

LMMs exhibited a lower statistical power (in particular for true associations located on the subcortical nuclei (**SFigure 2**). However, this result must be interpreted with caution as it may be due, in part, to the more stringent control of false positives, which means that overall fewer vertices reach significance (**Figure 2, STable 2)**. Yet, some of the power reduction may arise from the double fitting of the feature as fixed and random effect (Listgarten et al., 2012; Yang et al., 2014). A workaround (Yang et al., 2014) is to exclude the candidate vertex (and vertices strongly correlated) from the BRM calculation (Listgarten et al., 2012), though this requires computation of the BRM p times (complexity is O(pN^3^), with N the sample size and p the number of vertices), which becomes impractical for large sample sizes (Listgarten et al., 2012; Yang et al., 2014). In comparison, the current LMM implementation makes our analysis scalable to samples sizes of tens of thousands (computational complexity of O(pN^2^+N^3^+pN)) (Zhang et al., 2019). It should be noted that Restricted Maximum Likelihood (REML) estimation approach used in LMMs requires substantially more computational resources than the GLMs and thus requires the use of high performance clusters.

Beyond power and false positive rate, we observed from simulations that LMMs could pinpoint the grey-matter association with greater precision (smaller clusters of true positives, **Figure 3**). Lastly, we found that prediction achieved from clusters reaching significance in LMMs was on par with that from the best GLMs (**Figure 3**), despite fewer vertices included in the predictor. This suggests the higher specificity of LMMs. Overall, our simulations indicate that LMM with a single random effect currently offers a good trade-off between power and false positive rate. However, it still fails to ensure a cluster FWER below 5% (also reported on MWAS (Zhang et al., 2019)), despite a stringent Bonferroni correction to account for multiple testing.

Next, we applied the mass-univariate vertex-wise models to five real phenotypes of the UKB: age, sex, BMI, smoking status and fluid IQ. As in the simulations, the LMMs identified fewer vertices and clusters than the GLMs (**Figure 4, STable 2**). The LMM with multiple random-effect components was the most stringent (a single cluster of association), consistent with simulations which showed it had the lowest FWER and statistical power. In contrast, the LMM with a single random-effect component identified several cortical and subcortical associations with BMI, age and sex (**STable 3**). Most (12/19) of the top vertices (in each cluster) replicated in the UKB left out sample (**STable 3**), even though we cannot rule out that the same confounders might act similarly on the two UKB data sets. Overall, replication may be warranted to conclude about an association in future studies, considering the inflation of false positives (even when using LMMs, **Figure 2**). The top associated vertices with age, sex and BMI each captured less than 1% of the phenotypic variance, suggesting that many more small associations are likely to account for the full morphometricity of the phenotypes (**STable 2**). Our results echo the warning against the risk of small associations being confounded (e.g. by artefacts) in big-data neuroimaging (Smith & Nichols, 2018), which was confirmed by a recent exploratory study of putative MRI confounders in the UKB (Alfaro-Almagro et al., 2020). Note that LMMs can reduce false positive associations caused by correlations across and within the different types of measurements (**Figure 2, 3**). Finally, unlike in our simulations (**Figure 3**), LMMs resulted in lower prediction accuracy than GLMs in the UKB left out sample (**Figure 4**). Nonetheless, prediction from LMMs generalised better in another independent sample (**Figure 4**), the OASIS3 dataset (LaMontagne et al., 2019). This suggests that LMMs result in a more robust estimator of the association between trait and vertices.

In the past years, many studies have been published on the association between greymatter structure and our phenotypes of interest (see **STable 4-8** for a selective review of publications). Our simulation and empirical results suggest that some of these studies could report a substantial number of false positive or redundant associations. Nevertheless, due to the limitations outlined below, it is unclear which of these studies suffer from this issue and to which extent.

Firstly, it has been shown in the omics literature, that power of LMM may be reduced for phenotypes strongly associated with the covariation between features (Lloyd-Jones et al., 2017; Yang et al., 2014). This is likely the case for age and sex as indicated by their strong association with the PCs calculated from vertex-wise data (**STable 2**). This may be an important limitation for phenotypes associated with a cascade of changes in grey-matter, for which LMM would be over conservative.

LMM assumes a normal distribution of random effects, which may not be realistic for all phenotypes studied. It is equivalent to assuming highly regionalised, specialised brain regions, each displaying a small association with the phenotype. Several models have been proposed to relax this hypothesis, for example, to include large/outlying associations as fixed effects (stepwise LMM(Segura et al., 2012)), break down the feature list into sets of small and large associations (data driven approach: MOMENT(Zhang et al., 2019)), or consider more complex distributions using Bayesian LMMs (Bayesian alphabet (Lloyd-Jones et al., 2017; Moser et al., 2015)). They remain to be evaluated in the context of vertex-wise analyses. More simulations are warranted, to study other trait architectures, different trait distributions (e.g. skewed, discrete) or to evaluate more sophisticated models. Our framework of simulation may be easily adapted for such investigations, and offers the advantage of estimation of statistical power as well as false positive rate, which are not often reported at the same time (Eklund, Nichols, & Knutsson, 2016; Noble, Scheinost, & Constable, 2020).

The nature of the grey-matter regions identified in our GLM analyses of real phenotypes (for which the truth is unknown) can be a matter of debate, which depends on the (also unknown) nature of the correlation between vertices. Two key scenarios can explain the correlation but the data currently available to us does not allow to differentiate between them. First, the correlation could be solely due to confounders (e.g. **Figure 1b**), in which case the distal associations are false positives. Second, the correlation between vertices could reflect dynamic brain pathways relevant to the trait of interest. In this case, one could describe the GLM associations not found using LMM as redundant rather than false positives. Since we cannot differentiate between these two important causes of between-vertex correlation, we chose to label LMM models as parsimonious, until we understand better the effect of confounders on the vertices correlation structure as well as the longitudinal changes in greymatter and their relationship with the phenotypes.

Finally, three additional limitations are worthy of note as they may limit the interpretation of mass-univariate vertex-wise analyses (compared to GWAS results). First, grey-matter associations may be both causes or consequences of the phenotype studied, unlike GWAS findings, which can impact how to consider redundant associations. At one end of the spectrum are phenotypes such as age for which the direction of the causality is obvious (nothing causes chronological age). When describing which parts of the brain are affected by aging, one may be interested in reporting all associations, including all indirect and redundant. Though, there is no guarantee that those brain regions correctly map the brain pathway of ageing as they might also reach significance due to confusion factors. On the other hand, for many other phenotypes, the direction of causality is unclear (e.g. smoking, BMI...) and one may prefer a more parsimonious and robust brain mapping. Second, greymatter vertices are semi-arbitrary features which may be defined and measured in different ways (e.g. different cortical meshes in FreeSurfer). For instance, the resolution of the cortical tessellation is arbitrary and thus so is the number of local vertices which are found to be significant. Our analyses cannot rule out that results might be dependent on a particular processing or vertex definition that we did not consider in our analyses (e.g. volume processing from SPM, coarser surface mesh). Furthermore, we did not consider all possible covariates in GLM analysis, focussing on the more commonly used in previous analyses (age, sex, ICV, **STable 4-8).** More work is needed to evaluate the extended set(s) of covariates which have been recently proposed, from a large-scale study of the UKB data (Alfaro-Almagro et al., 2020). Thirdly, mass-univariate results may depend on the study sample used, which raises the question of generalisability into to samples from different age or ethnic groups or with different MRI qualities for instance.

In summary, we found that results obtained using the current state-of-the-art models (GLMs) used in MRI-trait association analyses likely suffer from a large inflation of false positive or redundant associations, due to the unaccounted correlation between vertices. In contrast, LMMs allow to control for all vertices fitted as a random effect, which result in a more parsimonious, robust and conservative characterisation of the localised associations between a phenotype and grey-matter structure. However, LMM results should still be interpreted with caution as our simulations show that the false positive rate remains higher than the standard type 1 error of 5%, even after Bonferroni correction.

## 8. URLs

Summary-level results: http://cnsgenomics.com/data.html;

Code available upon publication at https://github.com/baptisteCD;

OSCA: http://cnsgenomics.com/software/osca/;

ENIGMA protocols: http://enigma.ini.usc.edu/protocols/imaging-protocols/

## 9. Author contributions

PMV, NRW, JY, OC and BCD designed the analyses. FZ and JY developed the OSCA software. KK and JS assisted BCD with the UKB phenotypic and imaging data, including download, formatting and curation. BCD downloaded and processed the UKB MRI images. BCD performed the analyses and wrote the manuscript. All the authors reviewed the manuscript.

## 1.Acknowledgements

## 1.1. Data and code availability statement

Data used in this manuscript are held and distributed by the OASIS and UKB teams. We will release upon publication the scripts used in image processing and analyses to facilitate replication and dissemination of the results (see **URLs**). We will also release, the summary statistics of mass-univariate analyses performed on the UKB phenotypes.

## 1.4. Patient consent statement and ethics approval

Informed consent was obtained from all UK Biobank participants. Procedures are controlled by a dedicated Ethics and Guidance Council (http://www.ukbiobank.ac.uk/ethics), with the Ethics and Governance Framework available at http://www.ukbiobank.ac.uk/wp-content/uploads/2011/05/EGF20082.pdf. IRB approval was also obtained from the North West Multi-centre Research Ethics Committee. This research has been conducted using the UK Biobank Resource under Application Number 12505.

All necessary patient/participant consent has been obtained and the appropriate institutional forms have been archived by the OASIS team.

## 1.5. Other acknowledgements

We used R (R Development Core Team, 2012) (v3.3.3) for analyses not performed using OSCA (Zhang et al., 2019) and for plots. We used the colour-blind friendly R palette *viridis (Garnier, 2018), ukbtools (Hanscombe, 2017)* to facilitate UKB phenotype manipulation, *Morpho* and *Rvcg (Schlager, 2017)* to identify clusters, *rgl(Adler & Murdoch, 2020)* to generate brain plots of associations. Other packages used include, *dplyr (Hadley. Wickham & Francois, 2015), readr (H. H. Wickham, J.; Francois, R., 2017), rmarkdown (Allaire, 2018), matrixStats (Bengtsson, 2019), RcolorBrewer (Neuwirth, 2014), gridExtra (Auguie, 2017), ggplot2(Hadley Wickham, 2009), png (Urbanek, 2013), epuRate (Holtz, 2020).*

We would like to thank Allan McRae, the Institute of Molecular Bioscience (IMB) and the Research Computing Centre (RCC) IT teams at the University of Queensland for their support with high performance computing, data handling, storage and processing.

