## Supplementary Materials for "A parsimonious model for mass-univariate vertex-wise analysis"

### Supplementary Information for: **A parsimonious model for mass-univariate vertex-wise analyses**

**This PDF file includes:**

Figs. S1 to S16
Tables S1 to S7

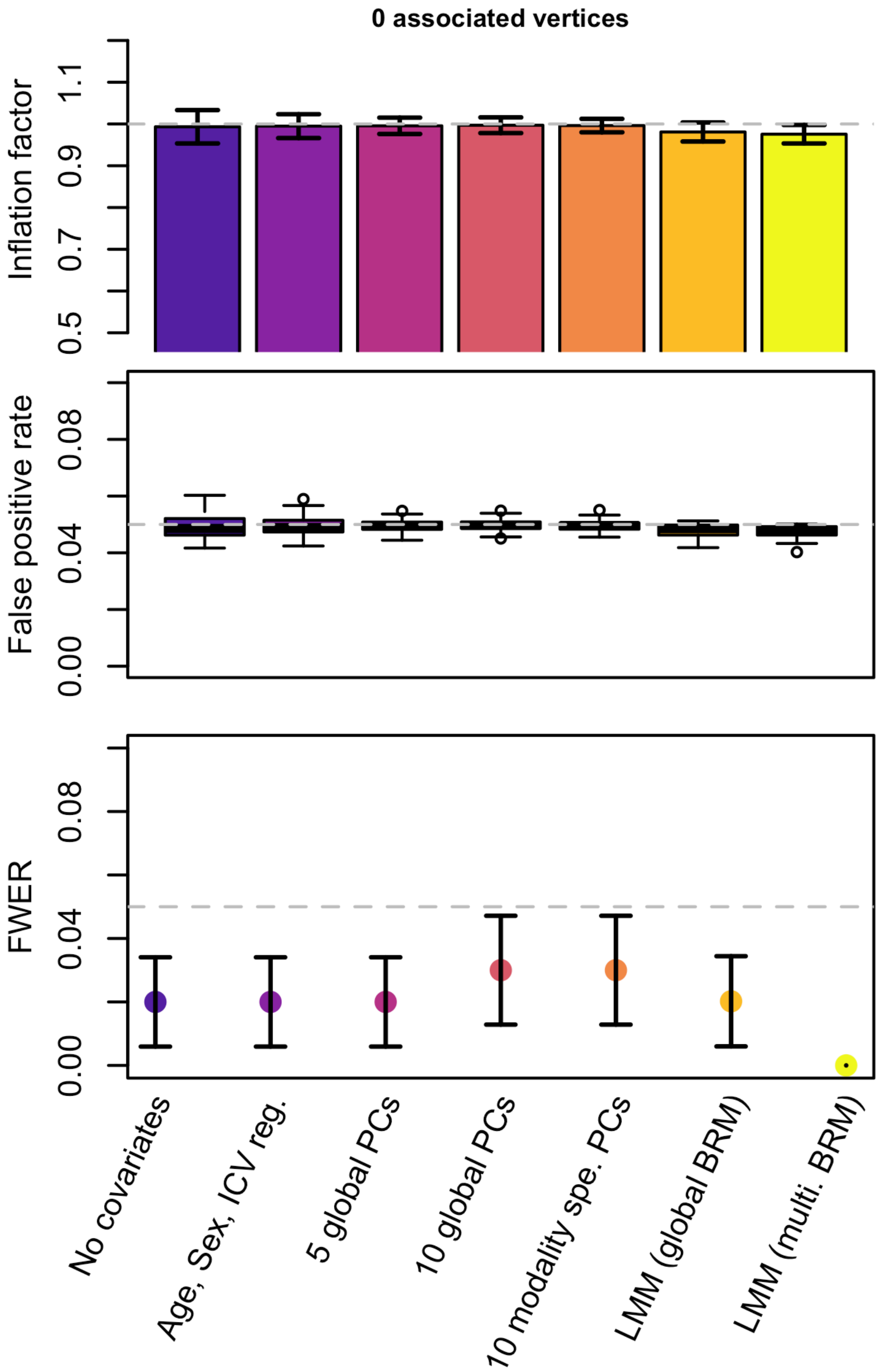


*SFigure 1: Summary of mass-univariate analyses on null phenotypes (not associated)*

A hundred phenotypes were simulated at random and analysed using each model.

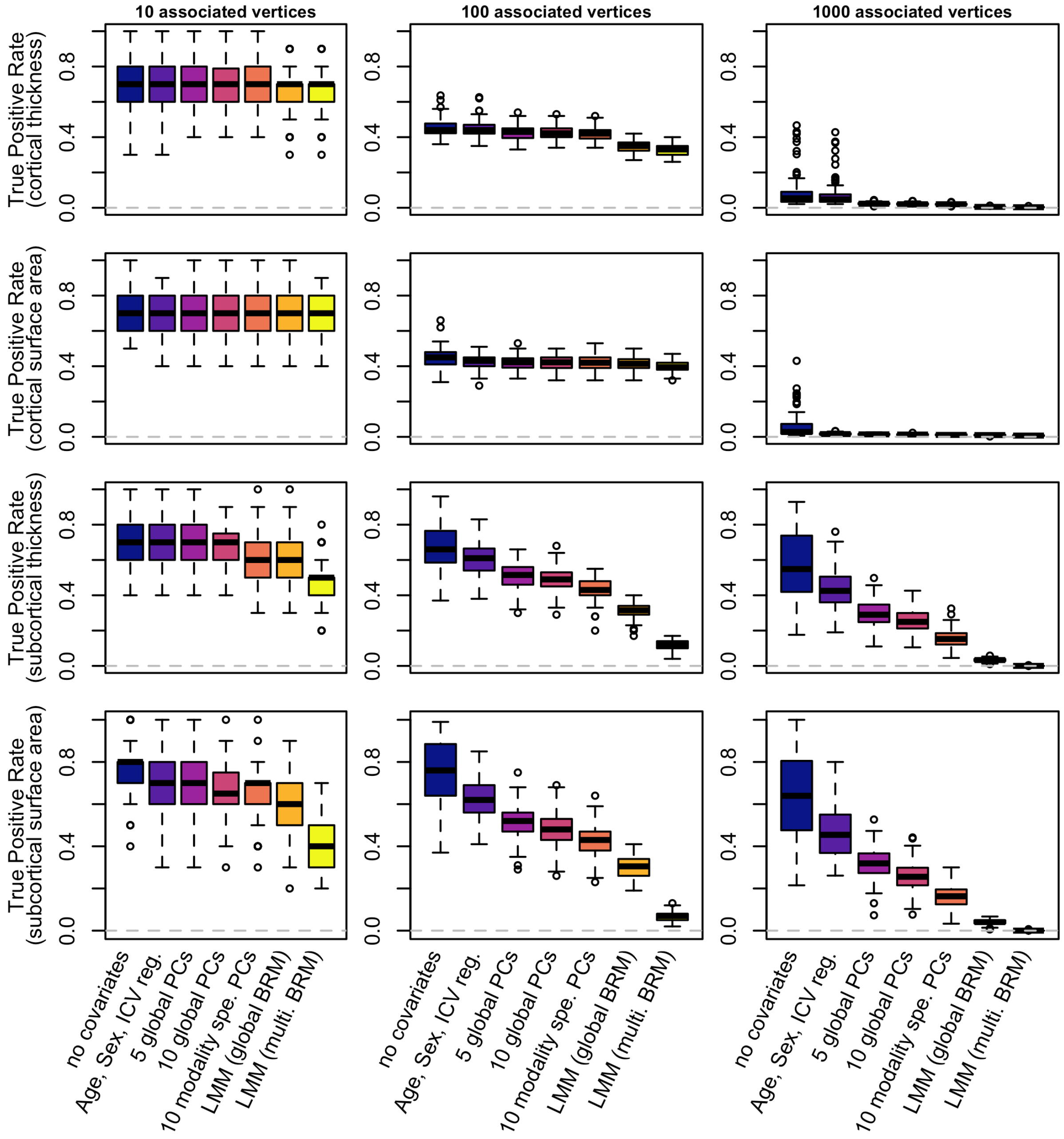


*SFigure 2: Statistical power for simulated phenotypes associated with a single type of modality*

*The phenotypes were simulated from a single type of measurement (indicated in parenthesis on the Y-axis label). The statistical power is measured using the true positive rate, or proportion of true associations significant after Bonferroni correction.*

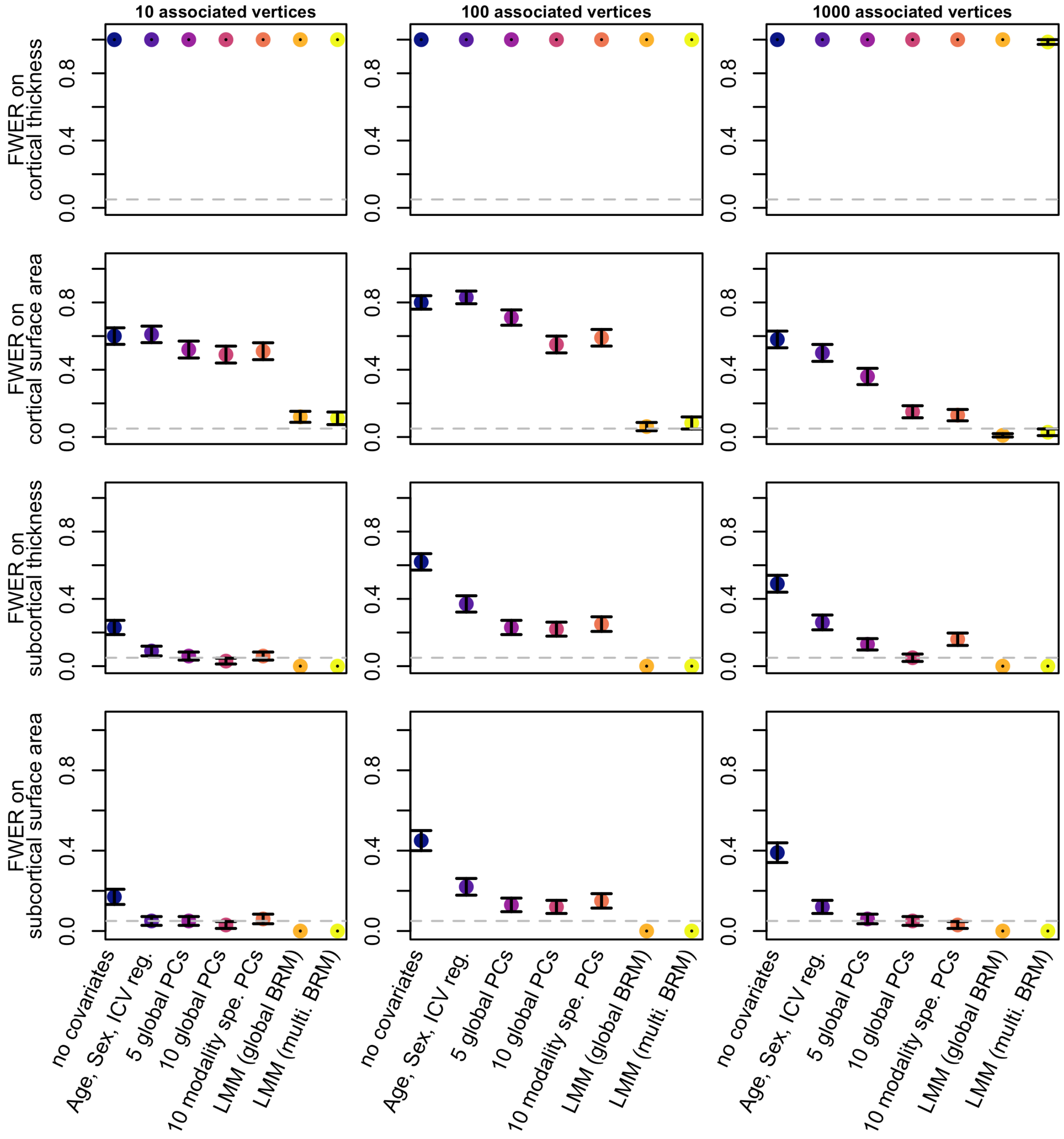


*SFigure 3: Probability of a false positive vertex arising from associations on cortical thickness*

*The phenotypes were simulated from cortical thickness only. The FWER corresponds to the proportion of replicates with a false positive vertex on each type of measurement.*

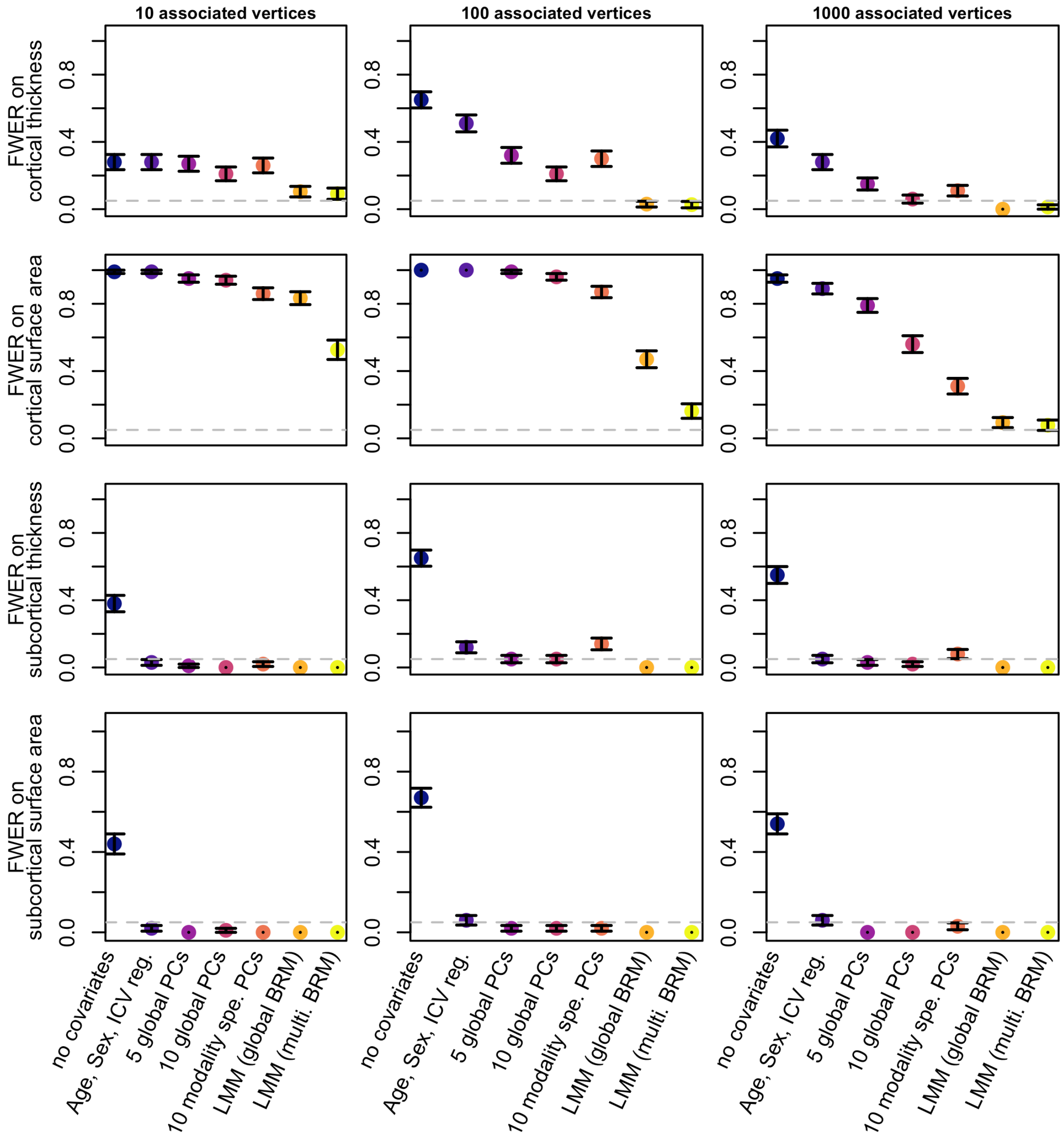


*SFigure 4: Probability of a false positive vertex arising from associations on cortical surface area*

*The phenotypes were simulated from cortical surface area only. The FWER corresponds to the proportion of replicates with a false positive vertex on each type of measurement.*

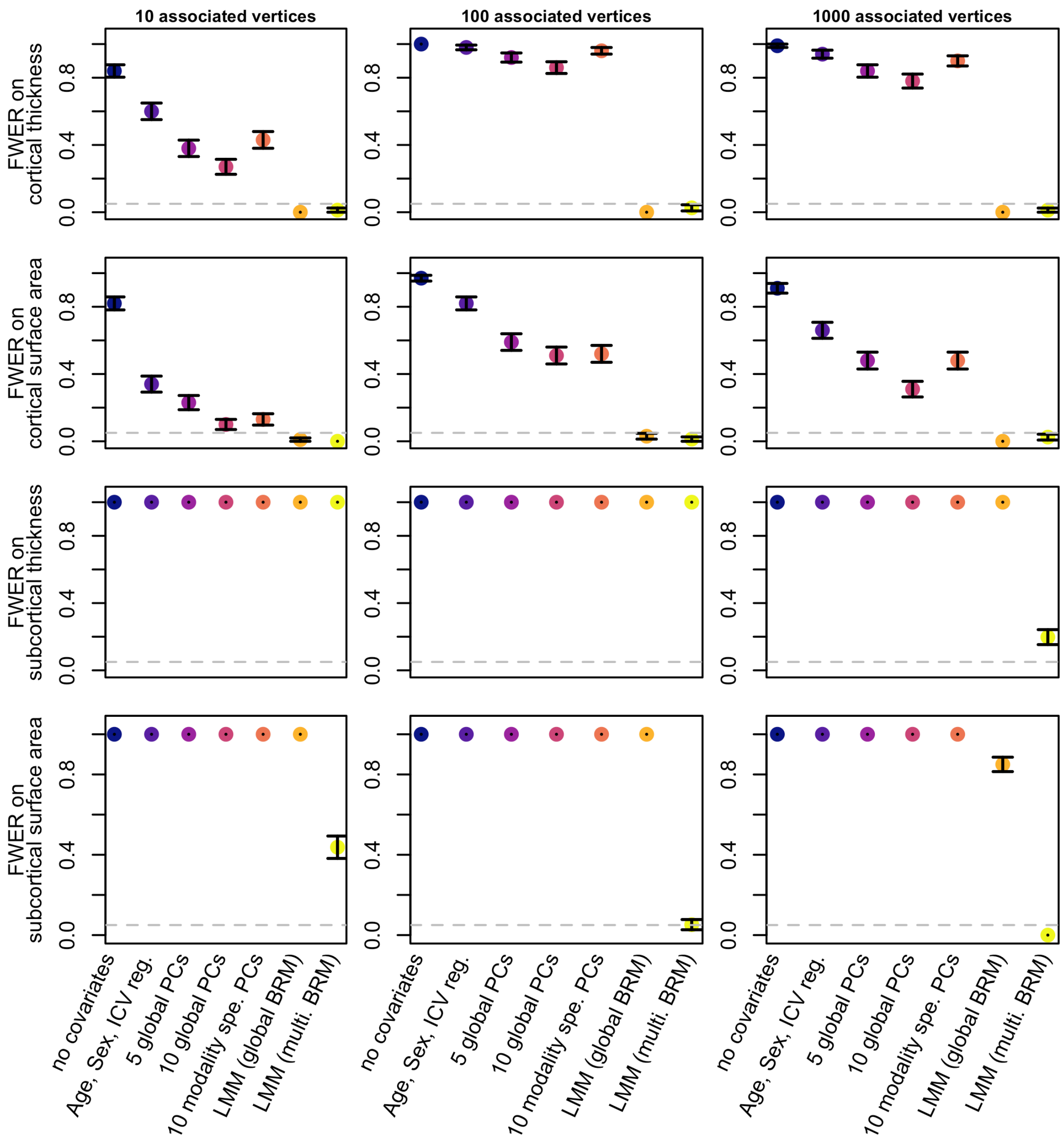


*SFigure 5: Probability of a false positive vertex arising from associations on subcortical thickness*

*The phenotypes were simulated from subcortical thickness only. The FWER corresponds to the proportion of replicates with a false positive vertex on each type of measurement.*

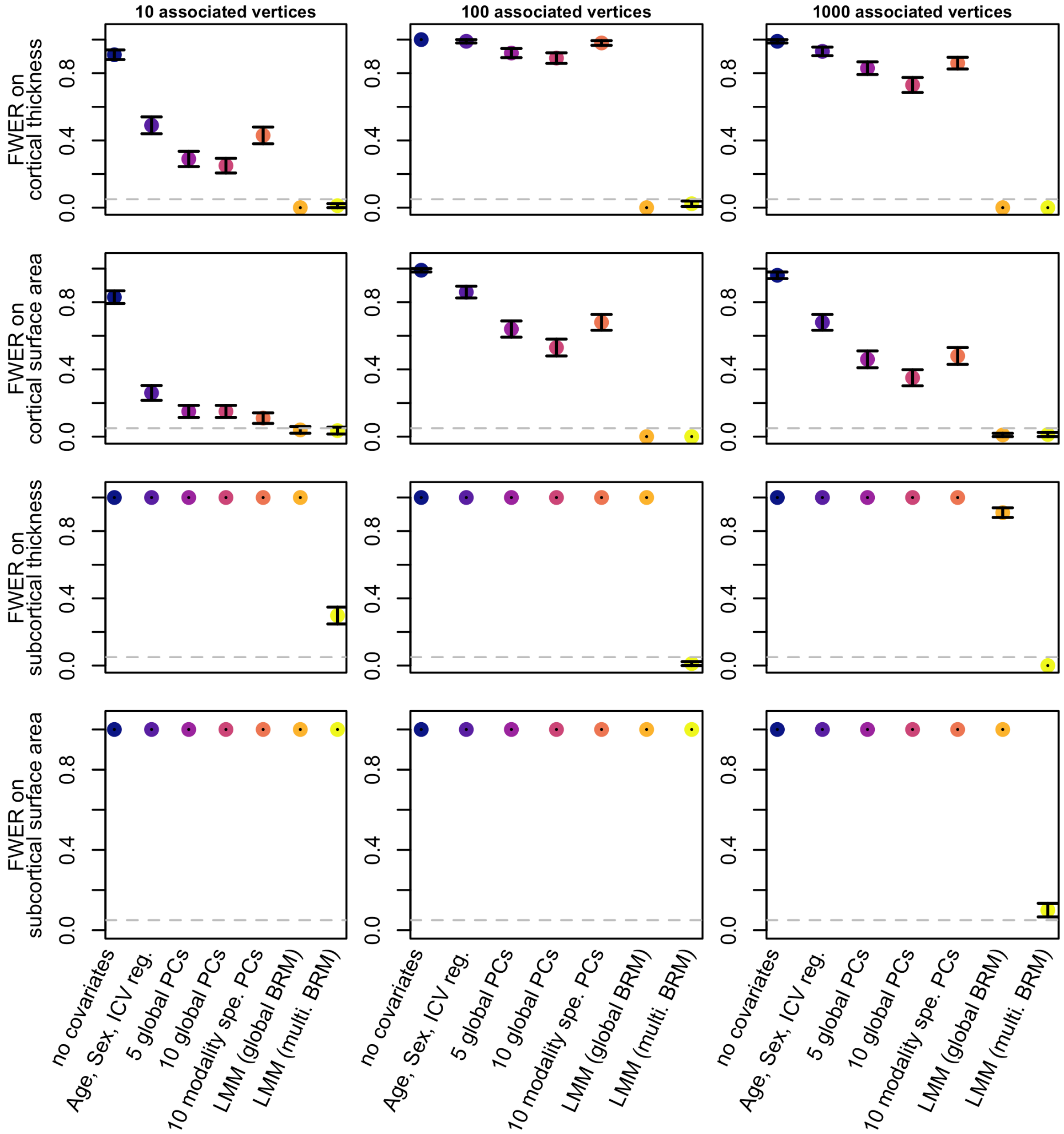


*SFigure 6: Probability of a false positive vertex arising from associations on subcortical surface area*

*The phenotypes were simulated from subcortical surface area only. The FWER corresponds to the proportion of replicates with a false positive vertex on each type of measurement.*

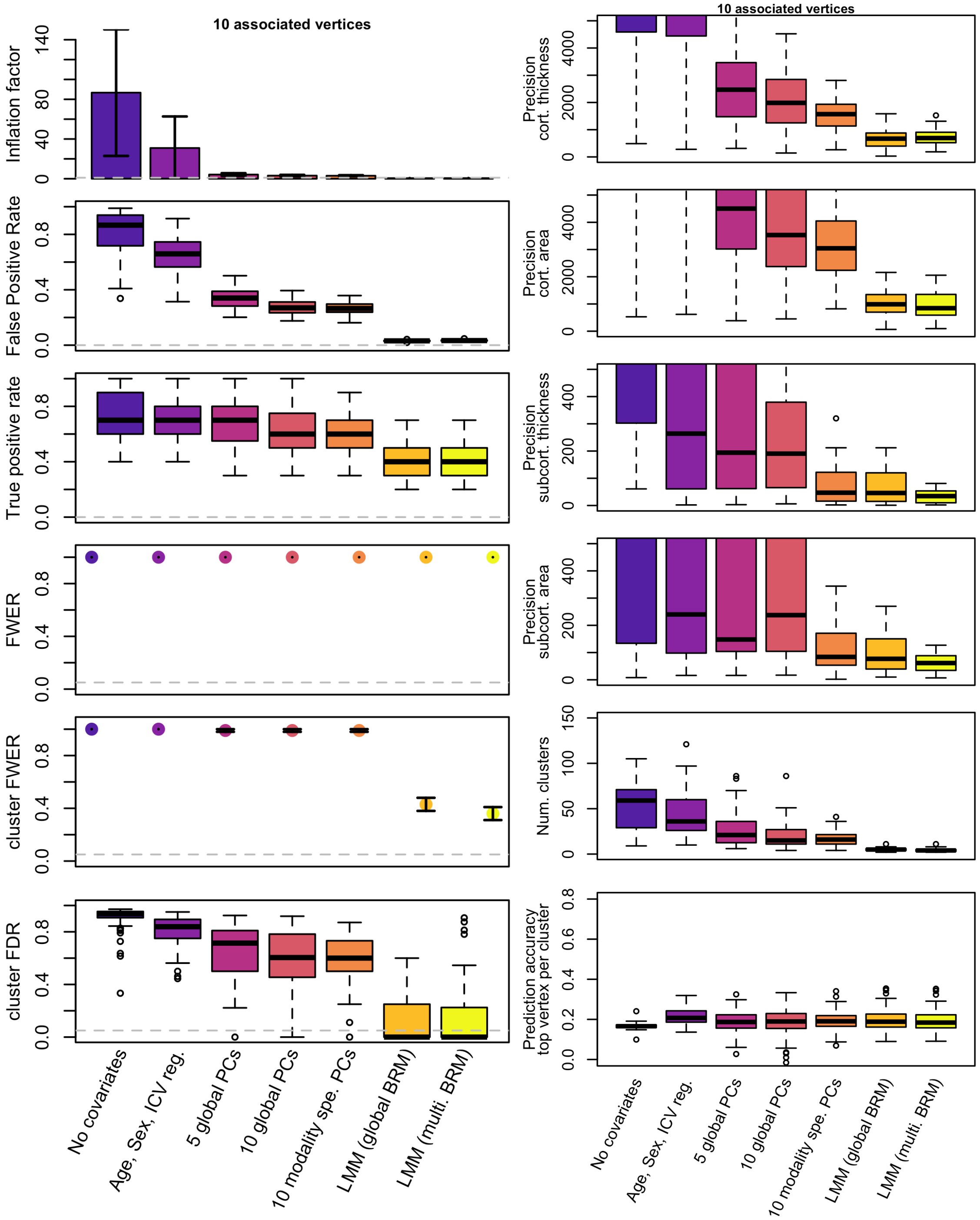


*SFigure 7: Results of mass-univariate analyses using smoothed meshes of cortical vertices.*

*For the ease of computation*, we only considered a simple simulation scenario of 10 associated vertices (morphometricity of 20%). Phenotypes were again simulated from the cortical and subcortical meshes, using the same seed vertices and weights as in previous simulations on unsmoothed data.

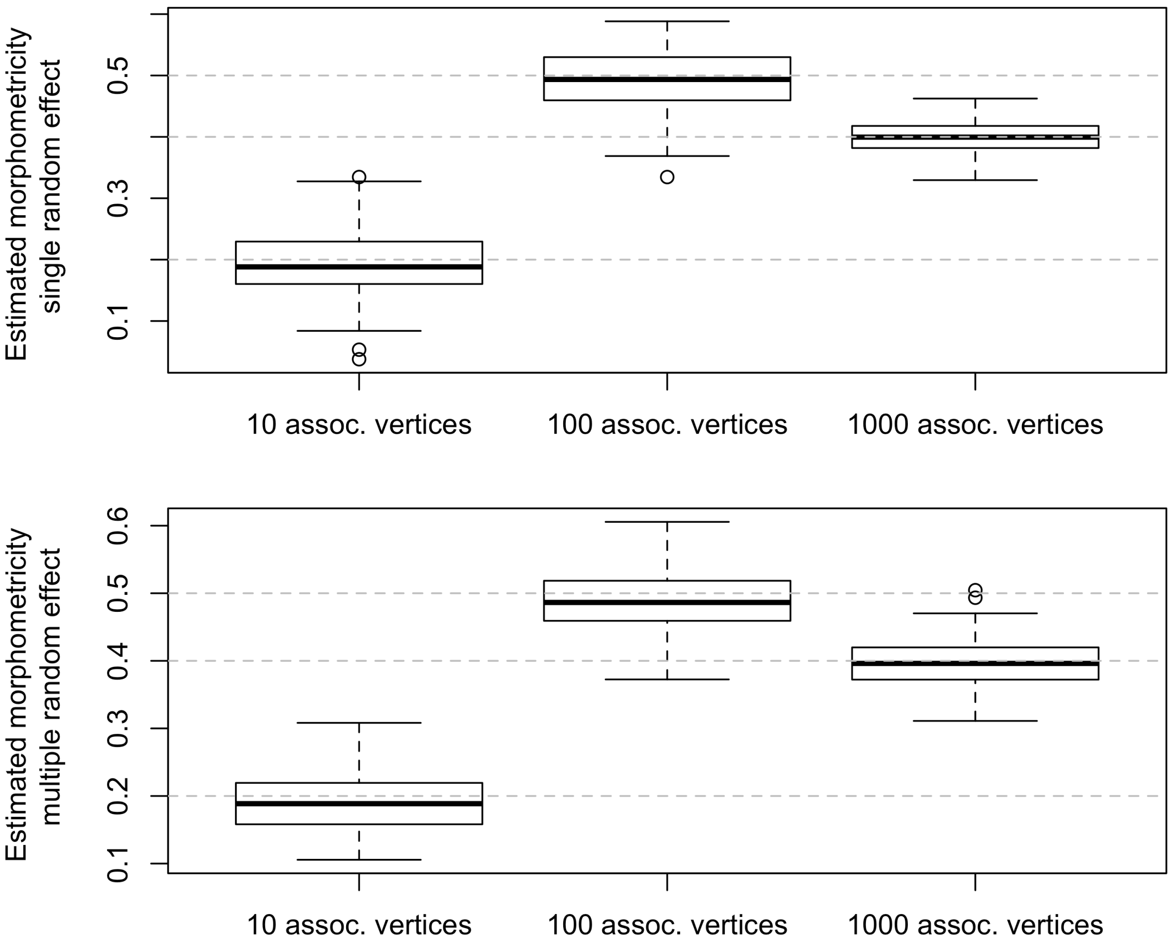


SFigure 8: Morphometricity estimates of our simulated traits from our two LMM models
The grey lines represent the expected values.

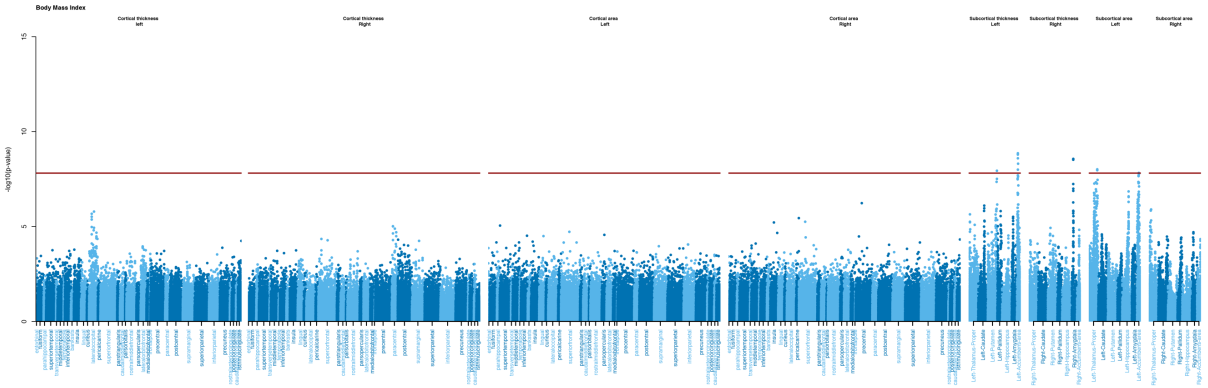


SFigure 9: Manhattan plot of mass-univariate vertex-wise analysis of body mass index
The Y axis shows the significance level (-log10 pvalue) for all grey matter vertices using the LMM “single random effect”. The red horizontal line indicates the brain-wide significance threshold using Bonferroni correction.

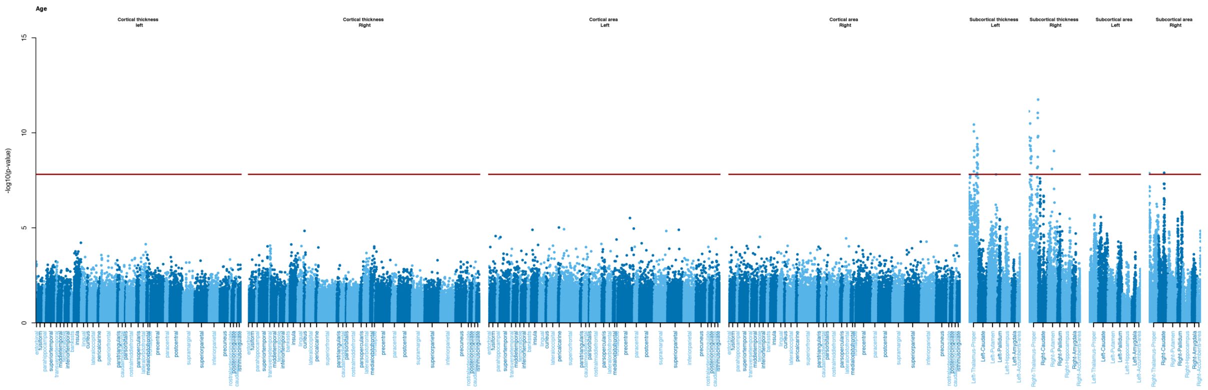

SFigure 10: Manhattan plot of mass-univariate vertex-wise analysis of age

The Y axis shows the significance level (-log10 pvalue) for all grey matter vertices using the LMM “single random effect”. The red horizontal line indicates the brain-wide significance threshold using Bonferroni correction.

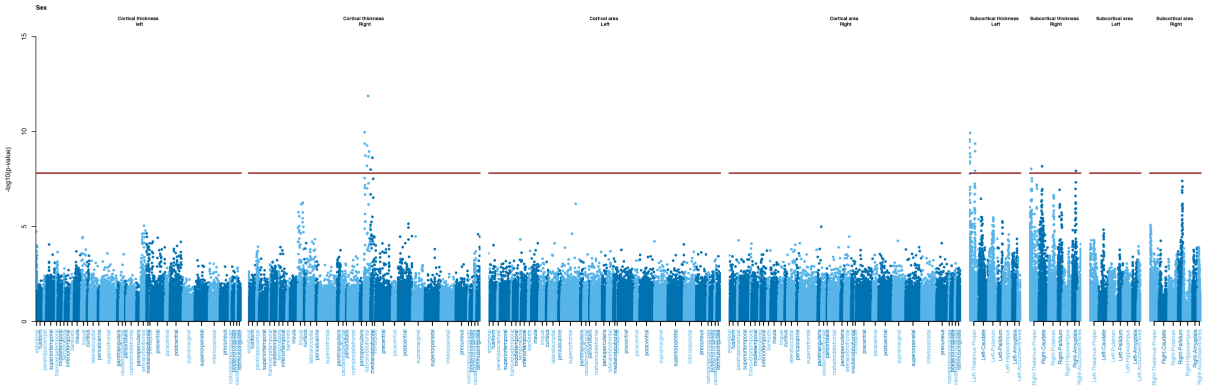


SFigure 11: Manhattan plot of LMM mass-univariate vertex-wise analysis of sex

The Y axis shows the significance level (-log10 pvalue) for all grey matter vertices using the LMM “single random effect”. The red horizontal line indicates the brain-wide significance threshold using Bonferroni correction.

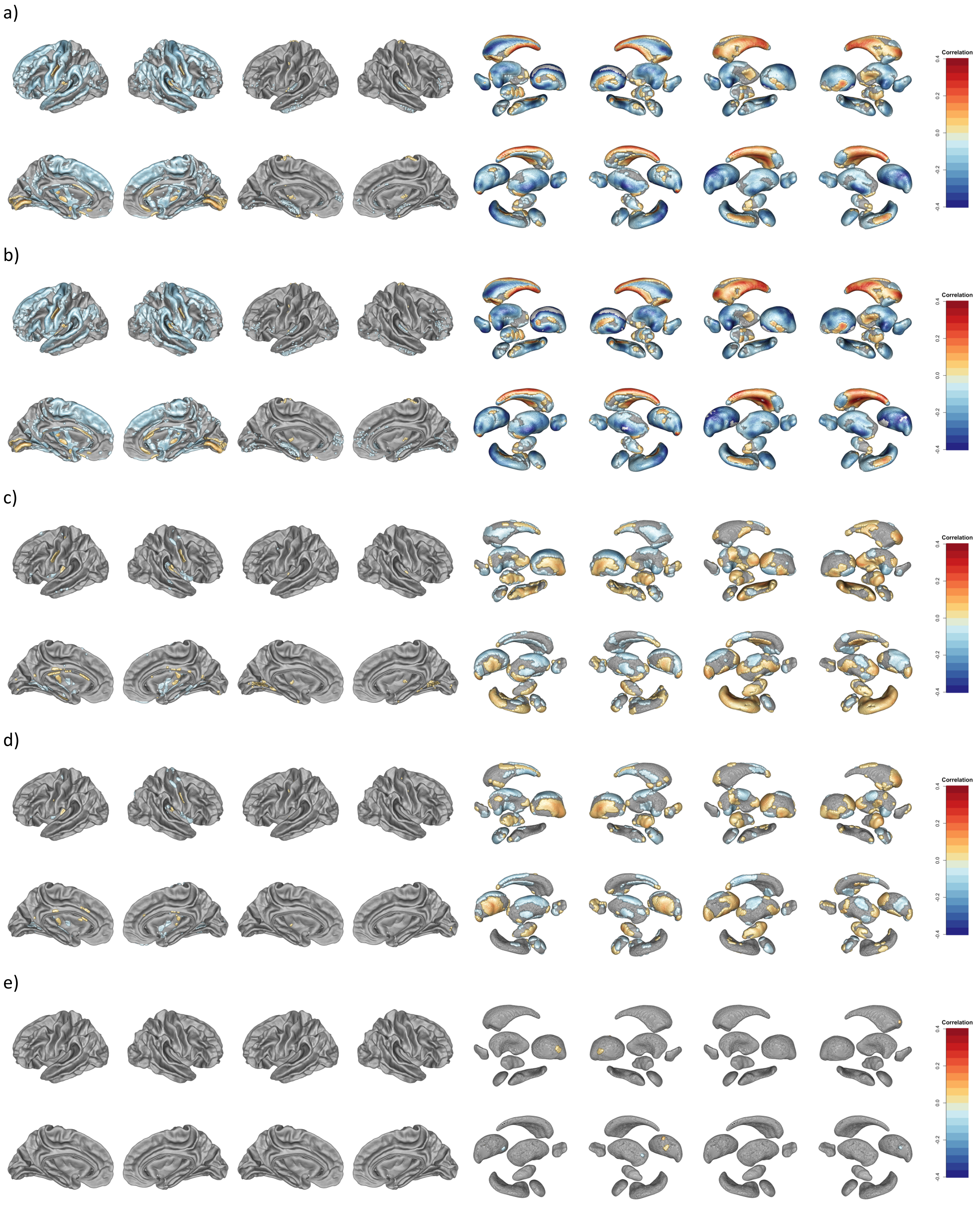


**SFigure 12: Results of vertex-wise analysis for age at MRI using the different models**

The brain plots present the significant vertices (in color) , for (from left to right) cortical thickness, cortical surface area, subcortical thickness and subcortical surface. The top and bottom rows shows the outside and inside view of the cortex and of the subcortical volumes. a) Results for GLM “no covariates”, b) GLM “sex, ICV”, c) GLM “5 global PCs”, d) GLM “10 global PCs, e) LMM “global BRM”.

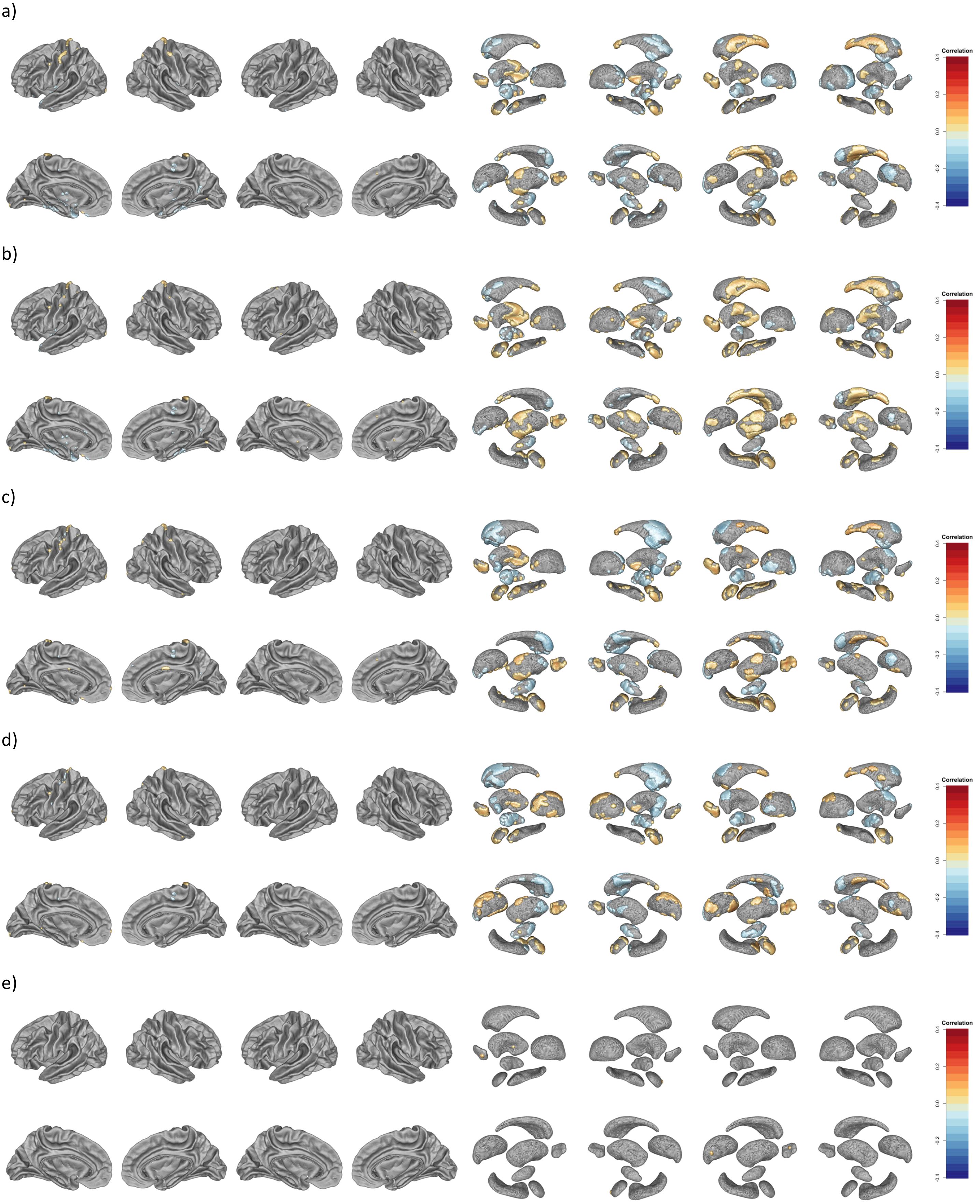


**SFigure 13: Results of vertex-wise analysis for BMI using the different models**

The brain plots present the significant vertices (in color) , for (from left to right) cortical thickness, cortical surface area, subcortical thickness and subcortical surface. The top and bottom rows shows the outside and inside view of the cortex and of the subcortical volumes. a) Results for GLM “no covariates”, b) GLM “sex, ICV”, c) GLM “5 global PCs”, d) GLM “10 global PCs, e) LMM “global BRM”.

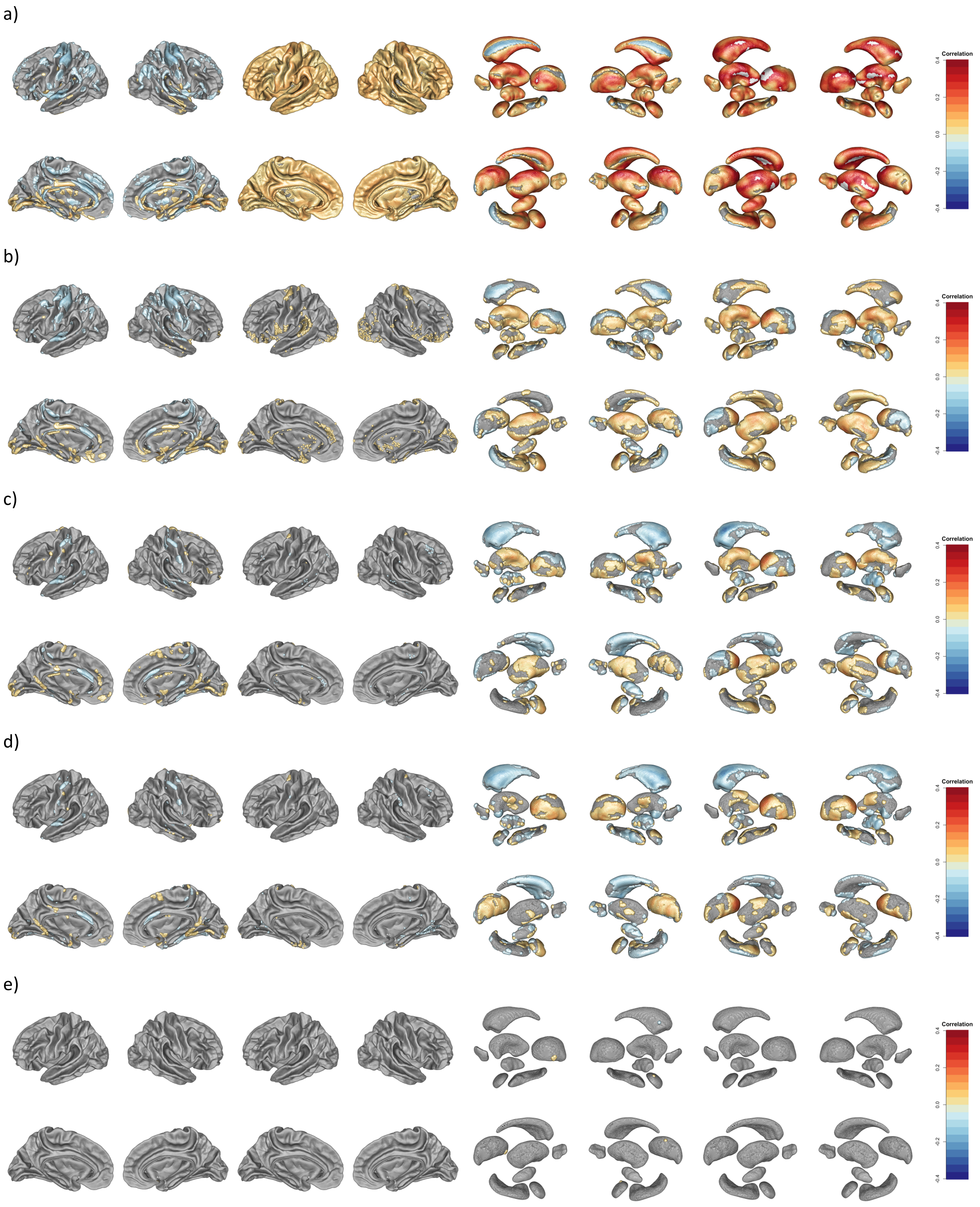


**SFigure 14: Results of vertex-wise analysis for sex using the different models**

The brain plots present the significant vertices (in color), for (from left to right) cortical thickness (left hemisphere, right hemisphere), cortical surface area, subcortical thickness and subcortical surface. The top and bottom rows shows the outside and inside view of the cortex and of the subcortical volumes. a) Results for GLM “no covariates”, b) GLM “sex, ICV”, c) GLM “5 global PCs”, d) GLM “10 global PCs, e) LMM “global BRM”.

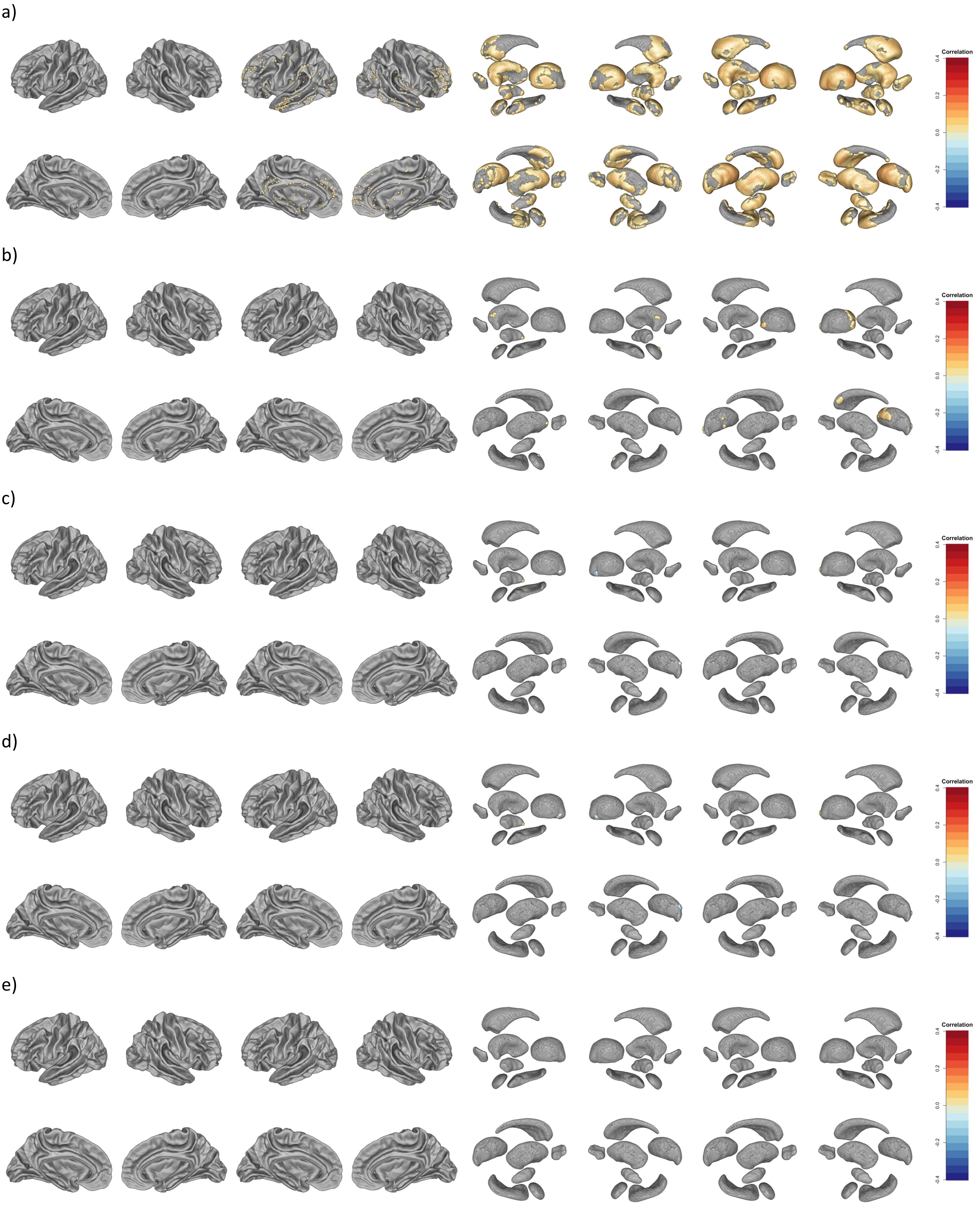


**SFigure 15: Results of vertex-wise analysis for fluid IQ using the different models**

The brain plots present the significant vertices (in color), for (from left to right) cortical thickness (left hemisphere, right hemisphere), cortical surface area, subcortical thickness and subcortical surface. The top and bottom rows shows the outside and inside view of the cortex and of the subcortical volumes. a) Results for GLM “no covariates”, b) GLM “sex, ICV”, c) GLM “5 global PCs”, d) GLM “10 global PCs, e) LMM “global BRM” (no significant association after multiple testing correction).

a)

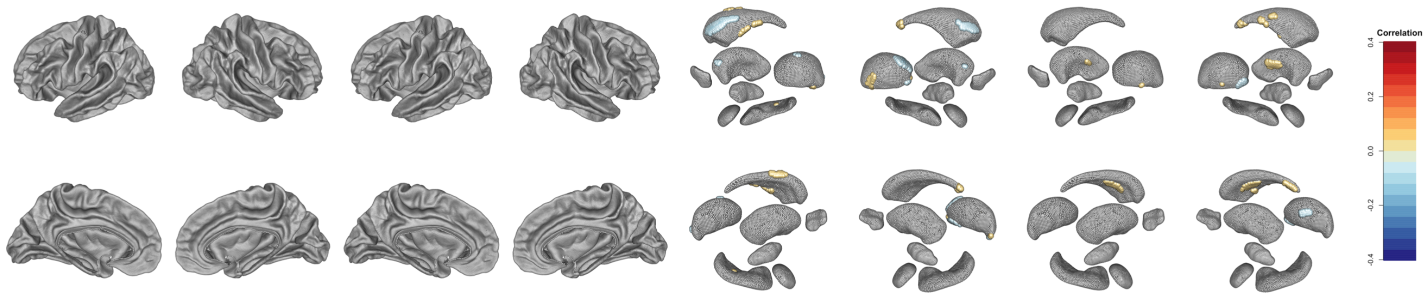


b)

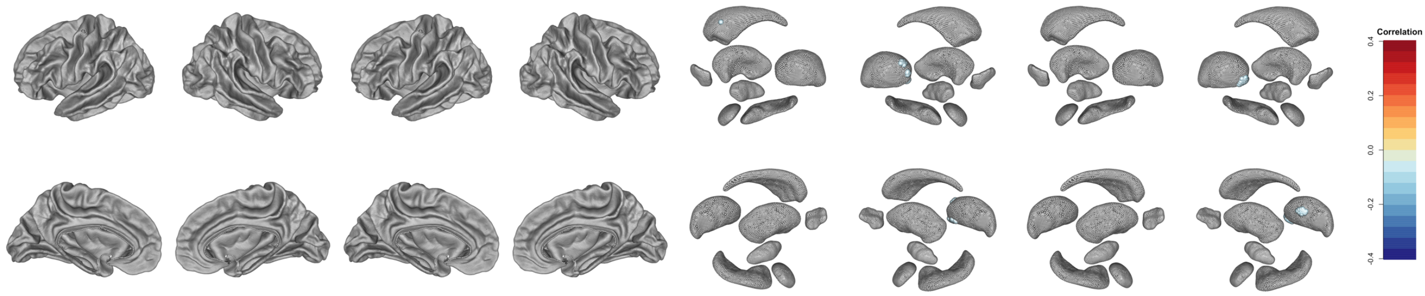


c)

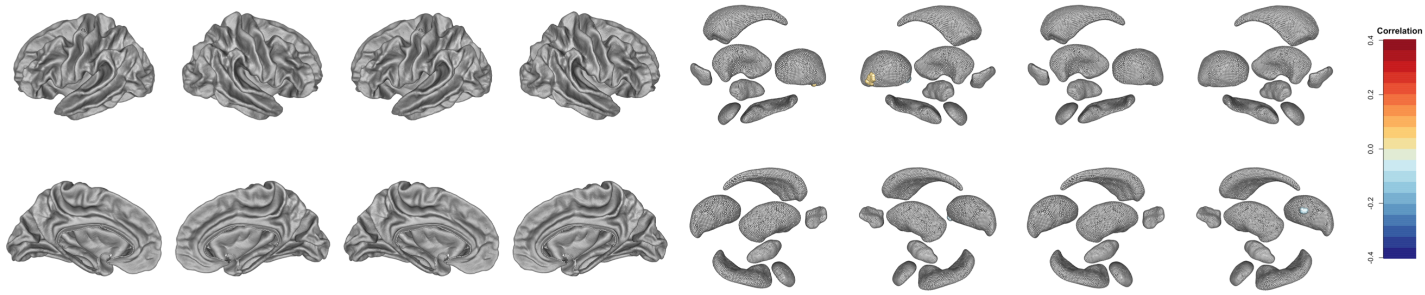


d)

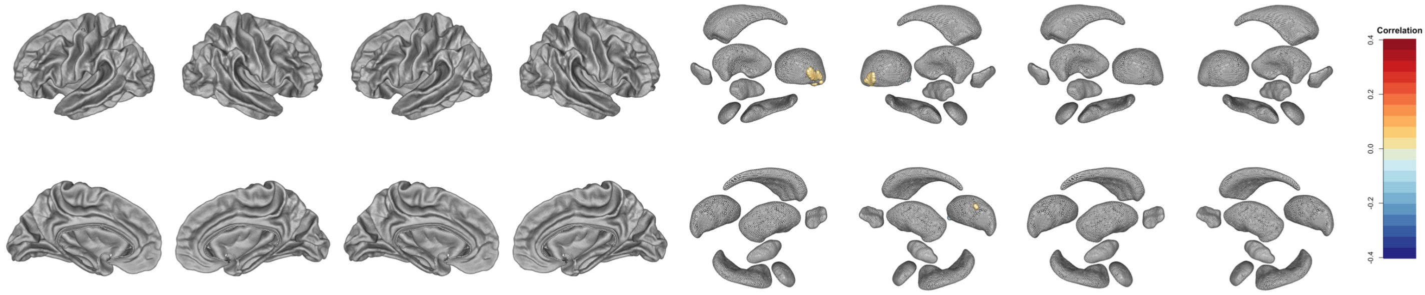


e)

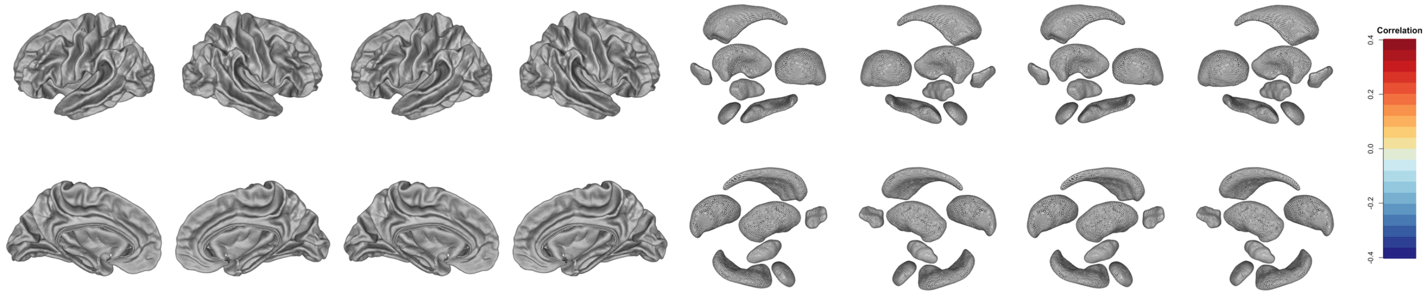


**SFigure 16: Results of vertex-wise analysis for smoking status using the different models**

The brain plots present the significant vertices (in color), for (from left to right) cortical thickness (left hemisphere, right hemisphere), cortical surface area, subcortical thickness and subcortical surface. The top and bottom rows shows the outside and inside view of the cortex and of the subcortical volumes. a) Results for GLM “no covariates”, b) GLM “sex, ICV”, c) GLM “5 global PCs”, d) GLM “10 global PCs, e) LMM “global BRM” (no significant association after multiple testing correction).

| Metric | Num assoc. vertices | **No covariates** | **Age, Sex, ICV reg.** | **5 global PCs** | **10 global PCs** | **10 modality spe. PCs** | **LMM (global BRM)** | **LMM (multi. BRM)** |
|---|---|---|---|---|---|---|---|---|
| **Lambda (inflation factor)** | 10 | 2.64 (1.58) | 1.63 (0.64) | 1.15 (0.07) | 1.12 (0.05) | 1.1 (0.04) | 0.94 (0.02) | 0.95 (0.02) |
| | 100 | 6.45 (4.27) | 3.01 (1.62) | 1.46 (0.09) | 1.37 (0.07) | 1.31 (0.05) | 0.91 (0.01) | 0.91 (0.01) |
| | 1000 | 4.42 (3.49) | 2.53 (1.08) | 1.48 (0.09) | 1.37 (0.05) | 1.32 (0.04) | 0.96 (0.01) | 0.97 (0.01) |
| **FPR (Nominal false positive rate)** | 10 | 0.2 (0.11) | 0.13 (0.06) | 0.07 (0.01) | 0.07 (0.01) | 0.07 (0.01) | 0.05 (<0.01) | 0.05 (<0.01) |
| | 100 | 0.39 (0.16) | 0.24 (0.1) | 0.12 (0.01) | 0.1 (0.01) | 0.1 (0.01) | 0.04 (<0.01) | 0.04 (<0.01) |
| | 1000 | 0.3 (0.14) | 0.21 (0.08) | 0.12 (0.01) | 0.1 (0.01) | 0.09 (0.01) | 0.05 (<0.01) | 0.05 (<0.01) |
| **True positive (TP) rate** | 10 | 0.72 (0.12) | 0.72 (0.12) | 0.71 (0.13) | 0.71 (0.13) | 0.7 (0.13) | 0.7 (0.13) | 0.69 (0.13) |
| | 100 | 0.45 (0.05) | 0.43 (0.04) | 0.42 (0.04) | 0.41 (0.04) | 0.41 (0.04) | 0.38 (0.04) | 0.37 (0.03) |
| | 1000 | 0.05 (0.04) | 0.03 (0.02) | 0.02 (<0.01) | 0.01 (<0.01) | 0.01 (<0.01) | 0.01 (<0.01) | 0.01 (<0.01) |
| **Median size of TP cluster on cortical surface area** | 10 | 11 (45) | 7 (32) | 3 (12) | 2 (9) | 1 (2) | 1 (1) | 1 (0) |
| | 100 | 1 (0) | 1 (0) | 1 (0) | 1 (0) | 1 (0) | 1 (0) | 1 (0) |
| | 1000 | 1 (0) | 1 (0) | 1 (0) | 1 (0) | 1 (0) | 1 (0) | 1 (0) |
| **Max. size** of TP cluster on cortical surface area | 10 | 63 (183) | 29 (79) | 18 (49) | 12 (37) | 7 (22) | 3 (8) | 2 (3) |
| | 100 | 163 (601) | 7 (17) | 3 (7) | 3 (5) | 2 (2) | 1 (1) | 1 (0) |
| | 1000 | 205 (1283) | 1 (2) | 3 (19) | 1 (0) | 1 (0) | 1 (0) | 1 (0) |
| **Median size of TP cluster on cortical thickness** | 10 | 152 (162) | 149 (168) | 124 (126) | 112 (77) | 107 (67) | 79 (45) | 77 (44) |
| | 100 | 81 (63) | 85 (172) | 50 (12) | 48 (13) | 45 (13) | 24 (8) | 24 (7) |
| | 1000 | 30 (22) | 26 (16) | 13 (7) | 13 (8) | 13 (7) | 7 (5) | 6 (5) |
| **Max. size** of TP cluster on cortical thickness | 10 | 384 (502) | 332 (304) | 254 (178) | 231 (142) | 218 (130) | 143 (65) | 139 (62) |
| | 100 | 1972 (8124) | 1240 (5624) | 226 (123) | 214 (107) | 196 (84) | 86 (20) | 85 (20) |
| | 1000 | 225 (325) | 196 (266) | 73 (94) | 50 (35) | 44 (33) | 11 (7) | 10 (8) |
| **Median size of TP cluster sub-cortical surface** | 10 | 795 (726) | 447 (488) | 236 (259) | 164 (152) | 104 (88) | 93 (80) | 49 (31) |
| | 100 | 855 (769) | 335 (434) | 127 (132) | 102 (99) | 57 (39) | 32 (22) | 25 (17) |
| | 1000 | 455 (551) | 140 (205) | 46 (57) | 41 (49) | 21 (17) | 6 (2) | 2 (NA) |
| **Max. size** of TP cluster sub- cortical surface | 10 | 804 (732) | 450 (486) | 239 (257) | 166 (151) | 106 (87) | 94 (79) | 50 (31) |
| | 100 | 1183 (889) | 568 (608) | 186 (210) | 150 (131) | 77 (57) | 41 (30) | 29 (18) |
| | 1000 | 813 (846) | 241 (370) | 53 (63) | 52 (55) | 23 (18) | 6 (2) | 3 (NA) |
| **Median size of TP cluster on sub-cortical thickness** | 10 | 747 (717) | 384 (479) | 206 (258) | 193 (236) | 82 (84) | 64 (62) | 32 (21) |
| | 100 | 417 (495) | 222 (249) | 90 (101) | 75 (81) | 37 (36) | 20 (19) | 15 (11) |
| | 1000 | 200 (220) | 116 (270) | 39 (44) | 30 (30) | 18 (30) | 7 (10) | 2 (2) |
| **Max. size** TP cluster on subcortical thickness | 10 | 759 (713) | 406 (481) | 212 (257) | 200 (234) | 85 (83) | 66 (63) | 32 (21) |
| | 100 | 668 (776) | 385 (494) | 148 (191) | 116 (131) | 51 (48) | 26 (28) | 17 (12) |
| | 1000 | 554 (673) | 188 (313) | 50 (61) | 39 (43) | 21 (31) | 7 (10) | 2 (2) |
| **FWER** | 10 | 1 (0) | 1 (0) | 1 (0) | 1 (0) | 1 (0) | 1 (0) | 1 (0) |
| | 100 | 1 (0) | 1 (0) | 1 (0) | 1 (0) | 1 (0) | 1 (0) | 1 (0) |
| | 1000 | 1 (0) | 1 (0) | 1 (0) | 1 (0) | 1 (0) | 0.97 (0.17) | 0.97 (0.17) |
| **Cluster FWER** | 10 | 0.99 (0.01) | 0.97 (0.02) | 0.98 (0.01) | 0.96 (0.02) | 0.85 (0.04) | 0.63 (0.05) | 0.49 (0.05) |
| | 100 | 1 (0) | 1 (0) | 1 (0) | 0.99 (0.01) | 0.97 (0.02) | 0.72 (0.05) | 0.72 (0.05) |
| | 1000 | 1 (0) | 1 (0) | 1 (0) | 0.99 (0.01) | 1 (0) | 0.7 (0.05) | 0.58 (0.05) |
| **Cluster FDR** | 10 | 0.75 (0.24) | 0.64 (0.24) | 0.48 (0.25) | 0.41 (0.24) | 0.32 (0.22) | 0.16 (0.17) | 0.12 (0.15) |
| | 100 | 0.78 (0.18) | 0.62 (0.17) | 0.33 (0.16) | 0.24 (0.13) | 0.17 (0.1) | 0.04 (0.03) | 0.03 (0.03) |
| | 1000 | 0.73 (0.13) | 0.65 (0.13) | 0.45 (0.16) | 0.34 (0.13) | 0.32 (0.13) | 0.13 (0.11) | 0.13 (0.15) |
| **Number of significant clusters** | 10 | 96 (125) | 33 (25) | 20 (17) | 15 (8) | 12 (5) | 9 (2) | 13 (19) |
| | 100 | 227 (204) | 135 (86) | 66 (21) | 55 (12) | 49 (8) | 39 (4) | 39 (7) |
| | 1000 | 142 (155) | 92 (77) | 30 (17) | 21 (7) | 18 (5) | 8 (3) | 7 (3) |
| **Prediction (r) from top vertex per cluster** | 10 | 0.29 (0.09) | 0.35 (0.07) | 0.38 (0.06) | 0.4 (0.05) | 0.42 (0.03) | 0.43 (0.02) | 0.43 (0.02) |
| | 100 | 0.32 (0.09) | 0.4 (0.08) | 0.54 (0.08) | 0.58 (0.06) | 0.61 (0.04) | 0.64 (0.02) | 0.63 (0.02) |
| | 1000 | 0.18 (0.04) | 0.2 (0.05) | 0.18 (0.05) | 0.18 (0.05) | 0.18 (0.04) | 0.15 (0.04) | 0.14 (0.04) |
| **Number of significant vertices** | 10 | 4636 (7293) | 1452 (1767) | 747 (552) | 616 (352) | 492 (230) | 325 (118) | 261 (110) |
| | 100 | 10596 (15205) | 6840 (12859) | 1967 (881) | 1666 (595) | 1309 (325) | 565 (100) | 510 (114) |
| | 1000 | 5813 (13178) | 2179 (2755) | 345 (315) | 209 (148) | 150 (83) | 23 (15) | 20 (15) |
| **Prediction (r) from significant vertices** | 10 | 0.22 (0.06) | 0.24 (0.05) | 0.25 (0.05) | 0.26 (0.06) | 0.26 (0.05) | 0.28 (0.06) | 0.28 (0.06) |
| | 100 | 0.23 (0.06) | 0.27 (0.06) | 0.34 (0.07) | 0.36 (0.06) | 0.38 (0.05) | 0.4 (0.04) | 0.4 (0.04) |
| | 1000 | 0.15 (0.05) | 0.16 (0.06) | 0.12 (0.05) | 0.12 (0.05) | 0.12 (0.04) | 0.12 (0.04) | 0.11 (0.03) |

STable 1: metrics of performance (SE) of mass-univariate models on simulated traits

The values are calculated over 100 simulated phenotypic traits.

*STable 2: Summary of mass-univariate vertex-wise analyses for the UKB phenotypes considered*

| | | BMI | Fluid Intelligence | Age | Sex | Smoking status |
|---|---|---|---|---|---|---|
| | *Adj. $R^2$ with age, sex & ICV* | 0.011 | 0.030 | 0.012 | 0.31 | 0.0086 |
| | *Adj. $R^2$ with first 10 PCs* | 0.033 | 0.032 | 0.41 | 0.43 | 0.0092 |
| **Uncorrected GLM** | *N assoc. vertices* | 10,947 | 24,776 | 136,278 | 355,130 | 707 |
| | *N assoc. clusters* | 232 | 640 | 970 | 714 | 34 |
| | *Max cluster size* | 862 | 2,030 | 22,358 | 130,651 | 116 |
| | *Prediction (UKB)* | 0.35 [0.33,0.38] | 0.17 [0.14,0.2] | 0.6 [0.58,0.62] | 0.68 [0.66,0.7] | 0.12 [0.09,0.15[illegible] |
| | *Prediction (OASIS3)* | 0.19 [0.13,0.25] | 0.2 [0.14,0.26] | 0.7 [0.67,0.73] | 0.65 [0.61,0.69] | 0.03 [-0.03,0.0[illegible] |
| **Age, sex, ICV GLM** | *N assoc. vertices* | 10,240 | 195 | 129,700 | 321,154 | 88 |
| | *N assoc. clusters* | 237 | 15 | 1269 | 1154 | 6 |
| | *Max cluster size* | 494 | 112 | 19,450 | 2,955 | 37 |
| | *Prediction (UKB)* | 0.35 [0.32,0.38] | 0.18 [0.15,0.21] | 0.52 [0.5,0.54] | 0.73 [0.72,0.75] | 0.09 [0.06,0.12[illegible] |
| | *Prediction (OASIS3)* | 0.14 [0.08,0.21] | 0.21 [0.15,0.27] | 0.64 [0.6,0.68] | 0.71 [0.68,0.74] | 0.03 [-0.03,0.1[illegible] |
| **5 global PCs GLM** | *N assoc. vertices* | 10,407 | 24 | 28,166 | 31,640 | 39 |
| | *N assoc. clusters* | 201 | 5 | 385 | 499 | 4 |
| | *Max cluster size* | 680 | 10 | 1,875 | 1,618 | 25 |
| | *Prediction (UKB)* | 0.35 [0.33,0.38] | 0.16 [0.13,0.19] | 0.44 [0.42,0.47] | 0.62 [0.6,0.64] | 0.11 [0.08,0.14[illegible] |
| | *Prediction (OASIS3)* | 0.24 [0.18,0.3] | 0.17 [0.11,0.23] | 0.28 [0.22,0.34] | 0.5 [0.45,0.55] | 0.01 [-0.05,0.0[illegible] |
| **10 global PCs GLM** | *N assoc. vertices* | 8,548 | 30 | 16,746 | 26,894 | 91 |
| | *N assoc. clusters* | 174 | 5 | 297 | 492 | 6 |
| | *Max cluster size* | 518 | 9 | 894 | 1782 | 43 |
| | *Prediction (UKB)* | 0.34 [0.31,0.37] | 0.16 [0.13,0.19] | 0.44 [0.41,0.46] | 0.58 [0.56,0.6] | 0.08 [0.05,0.11[illegible] |
| | *Prediction (OASIS3)* | 0.17 [0.1,0.23] | 0.17 [0.11,0.23] | 0.17 [0.11,0.23] | 0.58 [0.54,0.62] | -0.02 [-0.08,0.[illegible] |
| **10 moda. Spe. PCs GLM** | *N assoc. vertices* | 4,227 | 75 | 13,367 | 18,869 | 17 |
| | *N assoc. clusters* | 161 | 4 | 404 | 515 | 3 |
| | *Max cluster size* | 346 | 37 | 616 | 1437 | 6 |
| | *Prediction (UKB)* | 0.34 [0.32,0.37] | 0.17 [0.13,0.2] | 0.68 [0.66,0.69] | 0.69 [0.67,0.71] | 0.13 [0.1,0.16] |
| | *Prediction (OASIS3)* | 0.08 [0.02,0.15] | 0.16 [0.1,0.22] | 0.57 [0.53,0.61] | 0.59 [0.55,0.63] | 0.06 [0,0.12] |
| **Single random effect LMM** | *N assoc. vertices* | 11 | 0 | 47 | 27 | 0 |
| | *N assoc. clusters* | 5 | 0 | 8 | 6 | 0 |
| | *Max cluster size* | 5 | -Inf | 15 | 11 | -Inf |
| | *Morphometricity (SE)* | 0.16 [0.13,0.19] | NA [NA,NA] | 0.4 [0.38,0.43] | 0.48 [0.45,0.5] | NA [NA,NA] |
| | *Prediction (UKB)* | 0.13 [0.07,0.2] | NA [NA,NA] | 0.37 [0.32,0.43] | 0.47 [0.42,0.52] | NA [NA,NA] |
| | *Prediction (OASIS3)* | 0 | 0 | 0 | 9 | 0 |
| **Multiple random effect LMM** | *N assoc. vertices* | 0 | 0 | 0 | 1 | 0 |
| | *N assoc. clusters* | -Inf | -Inf | -Inf | 9 | -Inf |
| | *Max cluster size* | NA [NA,NA] | NA [NA,NA] | NA [NA,NA] | 0.13 [0.1,0.16] | NA [NA,NA] |
| | *Morphometricity (SE)* | NA [NA,NA] | NA [NA,NA] | NA [NA,NA] | 0.12 [0.06,0.18] | NA [NA,NA] |
| | *Prediction (UKB)* | 10947 | 24776 | 136278 | 355130 | 707 |
| | *Prediction (OASIS3)* | 232 | 640 | 970 | 714 | 34 |

*In the "age, sex and ICV adjusted" GLM, we dropped the corresponding covariate when studying age and sex. All adjusted $R^2$ are significant (pvalue<1e-16) considering the large sample size. Significance corresponded to a Bonferroni significance threshold of 1.5e-8, which accounts for the number of vertices and traits analyses.*

*STable 3: top vertices in each significant cluster from LMM single random effect models*

| | Top vertex | | | | | Discovery | | | | | Replication | |
|---|---|---|---|---|---|---|---|---|---|---|---|---|
| | | **hemi** | **modality** | **region** | **X;Y;Z coordinates** | **beta** | **se** | **pvalue** | **r** | **Cluster size** | **beta** | **pvalue** |
| *BMI* | LogJacs_10_738 | Left | Subcort. surface | Thalamus-Proper | 148.5 ; 110.9 ; 105.1 | 0.363 | 0.063 | 9.7E-09 | 0.082 | 2 | 0.029 | 7.3e-01 |
| | LogJacs_26_484 | Left | Subcort. surface | Accumbens-area | 136.3 ; 109.1 ; 144.7 | 0.408 | 0.072 | 1.5E-8 | 0.092 | 1 | 0.139 | 1.3e-01 |
| | thick_12_879 | Left | Subcort. thickness | Putamen | 152.4 ; 111.8 ; 125.4 | 0.316 | 0.055 | 1.2E-8 | 0.072 | 1 | 0.22 | 3.6e-03 |
| | thick_26_291 | Left | Subcort. thickness | Accumbens-area | 132.3 ; 114 ; 141.6 | 0.38 | 0.063 | 1.4E-09 | 0.086 | 5 | 0.359 | 3.2e-05 * |
| | thick_54_1331 | Right | Subcort. thickness | Amygdala | 106.2 ; 131.8 ; 141.1 | 0.447 | 0.075 | 2.7E-09 | 0.101 | 2 | 0.229 | 1.8e-02 |
| *Age* | LogJacs_49_1110 | Right | Subcort. surface | Thalamus-Proper | 108.9 ; 105.4 ; 108.6 | -0.552 | 0.097 | 1.3E-8 | -0.074 | 2 | -0.388 | 2.4e-03 |
| | LogJacs_50_2347 | Right | Subcort. surface | Caudate | 119.8 ; 100.6 ; 151.3 | 0.64 | 0.11 | 1.3E-8 | 0.086 | 1 | 0.388 | 1.2e-02 |
| | thick_10_2231 | Left | Subcort. thickness | Thalamus-Proper | 139.5 ; 109.8 ; 125.6 | -0.623 | 0.094 | 3.7E-11 | -0.084 | 5 | -0.658 | 5.6e-07 * |
| | thick_10_732 | Left | Subcort. thickness | Thalamus-Proper | 130.3 ; 107.4 ; 104.7 | 0.581 | 0.091 | 1.9E-10 | 0.078 | 15 | 0.793 | 2.9e-10 * |
| | thick_49_954 | Right | Subcort. thickness | Thalamus-Proper | 126 ; 107 ; 107.3 | 0.618 | 0.088 | 1.8E-12 | 0.083 | 11 | 0.577 | 3.4e-07 * |
| | thick_49_1337 | Right | Subcort. thickness | Thalamus-Proper | 110.2 ; 104.4 ; 111.3 | 0.536 | 0.081 | 3.2E-11 | 0.072 | 9 | 0.517 | 2.5e-04 * |
| | thick_49_1652 | Right | Subcort. thickness | Thalamus-Proper | 113.4 ; 97.9 ; 115.5 | 0.625 | 0.10 | 1.7E-09 | 0.084 | 2 | 0.76 | 3.2e-10 * |
| | thick_51_243 | Right | Subcort. thickness | Putamen | 98.6 ; 109.8 ; 117.3 | -0.43 | 0.070 | 9.1E-10 | -0.058 | 2 | -0.348 | 5.6e-04 * |
| *Sex* | rht_46930 | Right | Cort. thickness | Lateral orbitofrontal | -14 ; 11.2 ; -14.8 | 0.031 | 0.0043 | 1.3E-12 | 0.061 | 10 | 0.022 | 2.7e-04 * |
| | thick_10_1216 | Left | Subcort. thickness | Thalamus-Proper | 128.5 ; 113.5 ; 109.3 | 0.038 | 0.0058 | 1.2E-10 | 0.076 | 11 | 0.034 | 3.5e-05 * |
| | thick_10_2415 | Left | Subcort. thickness | Thalamus-Proper | 136.1 ; 111.9 ; 127.4 | 0.029 | 0.0046 | 4.3E-10 | 0.058 | 3 | 0.028 | 2.3e-05 * |
| | thick_49_1483 | Right | Subcort. thickness | Thalamus-Proper | 113.3 ; 97.8 ; 113.2 | 0.037 | 0.0064 | 9.2E-09 | 0.074 | 1 | 0.027 | 2.3e-03 |
| | thick_50_1785 | Right | Subcort. thickness | Right-Caudate | 123.2 ; 98.6 ; 141.6 | -0.034 | 0.0059 | 6.7E-09 | -0.068 | 1 | -0.031 | 1.3e-04 * |
| | thick_54_597 | Right | Subcort. thickness | Amygdala | 111.4 ; 121.1 ; 131.8 | 0.026 | 0.0046 | 1.2E-8 | 0.053 | 1 | 0.031 | 1.4e-06 * |

**: significant in the replication sample after multiple testing correction (p<0.05/85=5.8e-4).*

**STable 4: review of mass-univariate analyses of Age**

| Article | DOI | Year | Phenotype | N | population | Matching / covariates | Vertex modalities | Smoothing / mesh | Multiple testing | Significant regions |
|---|---|---|---|---|---|---|---|---|---|---|
| Medic et al., | 10.1038/ijo.2016.42 | 2016 | Age | 202 | Healthy adults | Age, sex, scanner, BMI, hemisphere, global thickness, area) | Cortical thickness, area | 15mm | Cluster level using Monte Carlo simulations | 11 regions in cortical thickness |
| Harrison et al., | 10.1016/j.neurobiolaging.2018.03.024 | 2018 | Successful ageing (cognition) | 129 | Older adults (70+) | sex | Cortical thickness | NS | No correction | Impossible to conclude |
| Dotson et al., | 10.3389/fnagi.2015.00250 | 2016 | Age | 46 | Middle aged adults (51-81 years | Sex, education, ICV/CT | Cortical thickness, surface | 10mm | FDR | 9 cortical regions surface area |
| Ducharme et al., | 10.1016/j.neuroimage.2015.10.010 | 2016 | Age | 384 | HC 4-22 years old, longitudinal (753 scans) | Sex, scanner (TBV) | Cortical thickness | 20mm | RFT peak and clusters | Linear model good approx. for most vertices |
| Li et al., | 10.1093/cercor/bhs413 | 2013 | age | 73 | Imaged at birth and 2 years, longitudinal | NA | Asymetry in sulcal depth, surface area, curvature. Left right SA ratio | NA | sufstat | Regional asymetries found for several modalities |
| Hogstrom | 10.1093/cercor/bhs231 | 2012 | Age | 322 | Healthy adults (20-85) | Sex, Total WM volume | Cortical thickness, surface, gyrification | 30mm | FDR Genovese et al., 2002 | Surface area showed strong age-related decreases, particularly pronounced in dorsomedial prefrontal, lateral temporal, and fusiform cortices, independently of total white matter volume. |
| Hugues et al., | 10.1016/j.neuroimage.2012.07.043 | 2012 | Age | 86 | Healthy subjects (20-74) | none | Thalamus shape/expansion | NA | FDR (non-specified) | Most of thalamus vertices significant |
| Sowell et al., | 10.1523/JNEUROSCI.1798-04.2004 | 2004 | Age | 45 | Imaged twice, age 5-11. T-test between t1 and t2 (no mixed models). | none | Cortical thickness (Eikonal Fire Equation, not FreeSurfer) 65K vertices p.h. | 15mm | Permutation to estimate minimal area significant | 10 significant regions |
| Muftuler et al., | 10.1016/j.brainres.2011.05.018 | 2011 | Age | 126 | Normally developing children age 6-10 | none | Cortical thickness (FS) | Unclear | FDR (unspecified) | Many cortical regions significant |
| Reid | 10.1002/hbm.20994 | 2010 | Age | 503 | nondemented elderly individuals (50–85 years) with a history ofsymptomatic cerebral small vessel disease (SVD | sex | Cortical thickness (CIVET) 40,962 vertices | NA | RFT | Most of the cortical sheet showed significant decrease with Age, with the greatest effects apparent in the ventrolateral prefrontal cortex(BA45, BA46, and BA47), the primary and secondary audi-tory cortices (BA41, BA42), Wernicke's area (BA22), medialtemporal lobe (BA36, BA28, excluding the hippocampal formation and amygdala), and the primary visual cortex. |
| Salat et al., | 10.1093/cercor/bhh032 | 2004 | Age | 106 | non-demented participants ranging in age from 18 to 93 years Imaged several times with T1 averaged | sex | Cortical thickness (FS) | 22mm | none | Some regions likely significant after bonferroni correction. |
| Gogtay | 10.1073pnas.0402680101 | 2004 | Age | 13 | Healthy age 4-21. Up to 4 scans per subject (52 images) | none | Cortical GM density | 15mm | None | Results not interpretable |

| | | | | | | | | | | |
|---|---|---|---|---|---|---|---|---|---|---|
| | | | | | | | (Thompson et al., 2000) | | | |
| Gogtay | 10.1002/hipo.20193 | 2006 | Age | 31 | Healthy age 4-21. Up to 4 scans per subject (100 images) | Sex, TBV | Hippocampus thickness, manual tracing, 30,000 measurements | NA | none | Results not interpretable |
| Van Soelen | 10.1016/j.neuroimage.2011.11.044 | 2012 | Age | 113 | Healthy twins 9–13 Imaged twice with 3 years interval | Sex, handedness, scan interval | Cortical thickness (CLASP) 40 962 vertices per hemisphere | 20mm | FDR (Genovese et al.,) | Widespread significant regions |
| Tamnes | 10.1523/JNEUROSCI.3302-16.2017 | 2017 | Age | 85 | Healthy adolescents, up to 2 scans per individual, 170 images total. 4 samples | Sex, scanning interval | Cortical thickness, volume, surface (FS) | 15mm | Monte carlo simulations, clusters p<0.05 | Widespread significant regions |

#### Stable 5: Review of mass-univariate analyses of Sex

| Article | DOI | Year | Phenotype | N | population | Matching / covariates | Vertex/voxel modalities | Smoothing / mesh | Multiple testing | Significant regions |
|---|---|---|---|---|---|---|---|---|---|---|
| Lotze et al., | 10.1038/s41598-018-38239-2 | 2018 | Sex | 2,838 | Adults age 21-90 | TBV, IQR, age, years of education, nicotine intake, alcohol consumption, and body mass index (BMI) | VBM | 8mm | FWER (not specified), cluster size >10 voxels | 25 significant regions |
| Chen et al., | 10.1016/j.neuroimage.2007.03.063 | 2007 | Sex | 411 | Adults, 44-48 years | age, years of education, handedness, and total intracranial volume | Cortical volume (VBM) | 12mm | FWER (not specified), threshold p<0.001 | 15 significant regions |
| Ruigrok et al., | 10.1016/j.neubiorev.2013.12.004 | 2014 | Sex | <2186 | Meta-analysis all ages. Incl. Chen et al., 2007 | Depending on study | Cortical volume, density (VBM) | Depends on study | FDR | 22 regions associated. Meta-analysis relies on foci, not on full map of summary statistics. |
| Jiang et al., | 10.1371/journal.pone.0073932 | 2013 | Sex | 266 | Inflamattory Bowel Disease (90). 176 HC | Age, Total grey matter volume | Cortical thickness | 8mm | FDR (RFT) | 4 cortical regions |
| Li et al., | 10.1093/cercor/bhs413 | 2013 | sex | 73 | Imaged at birth and 2 years, longitudinal | age | Asymetry in sulcal depth, surface area, curvature | | sufstat | |
| Boulos et al., | 10.1371/journal. pone.0152983 | 2016 | Sex | 87 | Right handed females (14-19 years) and males (14-18, See Chumachenko et al.,) | Age, IQ | Cortical thickness | 10mm | p<0.005, monte carlo simulations: cluster >250 mm2 | No significant findings |
| Richie et al,. | 10.1093/cercor/bhy109 | 2018 | Sex | 5216 | UK Biobank | Age, ethnicity | Cortical thickness, surface area, volume (FS)<br>Rs-fMRI<br>WM microstructure | 20mm | None (post hoc analysis) | Results compared to ROI based results.<br>Tred: large SQ and VOL in males, larger thickness in females |
| Luders et al | 10.1002/hbm.20187 | 2005 | Sex | 60 | Healthy young adults, right handed, matched for age | 2 processing, one accounting for global head size in realignment | Cortical thickness (Eikonal fire equations) | 15mm | permutation | Increased CT in females, widespread. Esp. when accounting for head size |
| Lv et al., | 10.1016/j.neuroimage.2010.05.020 | 2010 | Sex | 184 | Healthy adults (18-70) | Age, head size (GM+WM+CSF) | Cortical thickness (no FS) 40,962 p.h. | 20mm | FDR (Genovese et al.,) | cortical thickening in females appeared extensively in the frontal, parietal and occipital lobes, including the superior frontal gyrus, precentral gyrus, and postcentralgyrus in both hemispheres, and the superior parietal lobule, cuneus, and frontal pole in left hemispheres. The male cortex was significantly thicker than that of the female only in some small regions of the temporal lobes. |
| Sowell et al., | 10.1093/cercor/bhl066 | 2007 | Sex | 176 | Healthy 7-87 years | Age, age2, (height) | Cortical thickness | | Permutations (within large | Results of ROI permutation analyses (shown in Table 2) confirm the significance of sex differences in |

| | | | | | | | | | | |
|---|---|---|---|---|---|---|---|---|---|---|
| | | | | | | | (manual and automated processing) Eikonal fire equation 65536 vertices p.h. | | cortical regions) to estimate cluster size | cortical thickness in right lateral parietal (P = 0.048), right lateral temporal (P= 0.024), and left medial occipital (P = 0.017) regions. |
| | | | | 36 | Follow up in male female sample matched for TBV | | | | | Female thicker. Right: Lateral ventral frontal<br>Lateral occipital<br>Lateral parietal<br>Lateral temporal |
| Van Velsen | 10.1016/j.neulet.2013.06.063 | 2013 | Sex | 1022 | Non-demented elderly: age ~68 | ICV, education (in men and women separately) | Cortical thickness (FS) | NA | none | Results not presented/discussed. ROI focus |
| Reid | 10.1002/hbm.20994 | 2010 | Sex | 503 | nondemented elderly individuals (50–85 years) with a history ofsymptomatic cerebral small vessel disease (SVD | age | Cortical thickness (CIVET) 40,962 vertices | NA | RFT | Vertex-wise analyses highlightsome regions where a moderate Sex effect was apparent.AfterP-value correction, these effects were not significant; |

**Stable 6: Review of mass-univariate analyses of smoking (and related traits)**

| Article | DOI | Year | Phenotype | N | population | Matching / covariates | Vertex modalities | Smoothing / mesh | Multiple testing | Significant regions |
|---|---|---|---|---|---|---|---|---|---|---|
| Jorgensen et al., | 10.1503/jpn.140163 | 2015 | Smoking | 743 | 237 healthy controls and 506 psychiatric cases | Age, sex, diagnosis | Cortical thickness | 20mm | FDR (no reference) | 1 cluster significant in patient sample |
| | | | Smoking amount | | | | | | | No significant result |
| Cox et al., | 10.1093/eurheartj/ehz100 | 2019 | Vascular risk factors (BMI, smoking) | 7928 | UKB adults | age, sex, ethnicity, head size (for volumetric data), and head positioning confounds | Cortical volume | 20mm | FDR (Benjamini Hochberg) | Several large cortical regions (lateral and medial temporal lobes) |
| Chye et al., | 10.1111/adb.12830 | 2019 | Substance dependence (incl. nicotine) | 3905 | Multiple substance dependence (ENIGMA) | Site, sex, age, and ICV | Subcortical thickness and area | none | FDR (Landers et al.,) | Several subcortical volumes associated |
| Boulos et al., | 10.1371/journal.pone.0152983 | 2016 | Substance abuse (10 substances, incl. tobacco) | 43 | Right handed females (14-19 years) | Age, IQ | Cortical thickness | 10mm | p<0.005, monte carlo simulations: cluster >250 mm2 | pregenual rostral anterior cingulate cortex extending to the medial orbitofrontal cortex |
| Chumachenko | 10.3109/00952990.2015.1058389 | 2015 | Substance use (and conduct problems) | 44 | Males 14-18 | Age, IQ, total cortical thickness | Cortical thickness | NA | FWER – cluster leve; - Monte Carlo simulation (10,000 iterations) with a cluster-forming threshold vertex-level p-value of 0.005 (55) | Left posterior cingulate/precuneus |

#### Stable 7: Review of mass-univariate analyses of BMI (or related traits)

| Article | DOI | Year | Phenotype | N | population | Matching / covariates | Vertex modalities | Smoothing / mesh | Multiple testing | Significant regions |
|---|---|---|---|---|---|---|---|---|---|---|
| Medic et al., | 10.1038/ijo.2016.42 | 2016 | BMI | 202 | Healthy adults | Age, sex, scanner, hemisphere, global thickness, area, (smoking status) | Cortical thickness, area | 15mm | Cluster level using Monte Carlo simulations | 2 clusters in cortical thickness |
| Sharkey et al., | 10.3389/fnins.2015.00024 | 2015 | BMI | 378 (716 scans) | Healthy children (<18), longitudinal MRIs | Age, sex, scanner | Cortical thickness (CIVET) | 20mm | FDR correction (Non-specified, surfstat?) | No significant association |
| Bernardes et al., | 10.1007/s11011-018-0223-5 | 2018 | BMI / obesity | 31 lean normoglycemic controls<br>44 obese | 28 Obese with T2Diabetes<br>Age 40-70 | Age, sex, hypertention and ICV | Cortical thickness, area, volume | 10mm | Monte-carlo simulations, Pthreshold <0.01 | 1 cortical thickness cluster |
| Veit et al., | 10.1016/j.nicl.2014.09.013 | 2014 | BMI | 72 | Healthy subjects age 19-50 | Age, sex, total surface area, education | Cortical thickness | 10mm | Monte carlo threshold estimation, after cutoff P<0.05 | 3 thickness clusters |
| Varma et al., | 10.1002/hipo.22586 | 2016 | Physical activity | 90 | Adults > 6o years | intracranial volume (ICV), age, years of education, body mass index (BMI), cardiovascular disease burden (CVD), and global cognitive function | Subcortical shape | NA | FWER and FDR | Some significant regions (hippocampus) |
| Cox et al., | 10.1093/eurheartj/ehz100 | 2019 | Vascular risk factors (BMI, smoking) | 7928 | UKB adults | age, sex, ethnicity, head size (for volumetric data), and head positioning confounds | Cortical volume | 20mm | FDR (Benjamini Hochberg) | Several large cortical regions (lateral and medial temporal lobes) |
| Leritz et al., | 10.1016/j.neuroimage.2010.10.050. | 2011 | Cerebrovascular health (PCs derived from BMI) | 115 | Healthy controls, age 43-83 | age | Cortical thickness | 20mm | Clustering, after P<0.05 | Comparison with other results impossible |

**Stable 7: Review of mass-univariate analyses of IQ/cognition**

| Article | DOI | Year | Phenotype | N | population | Matching / covariates | Vertex modalities | Smoothing / mesh | Multiple testing | Significant regions |
|---|---|---|---|---|---|---|---|---|---|---|
| Harrison et al., | 10.1016/j.neurobiolaging.2018.03.024 | 2018 | Successful ageing (cognition) | 129 | Older adults (70+) | sex | Cortical thickness | NS | No correction | Impossible to conclude |
| Abe et al., | 10.1111/acps.12922 | 2018 | Executive functioning | 160 | HC, Type I and II bipolar | Sex (no age) + lot more in sensitivity analyses | Cortical thickness | 10mm | Monte carlo, cluster wise. Threshold p<0.05 | Several regions, some found across disease groups |
| Navas-Sanchez | 10.1002/hbm.23143 | 2016 | Math gifted | 62 | Spanish adolescents – IQ matched | age, gender, and IQ | Cortical thickness, area, volume | 15mm | Cluster wise probability method (FDR) Hagler et al., 2006 | Surface and thickness associated regions |
| Burgaleta | 10.1002/hbm.22305 | 2014 | Fluid IQ (and other IQ dimensions) | 104 | Psychology undergraduates (age ~19) | Age, sex (brain size in processing) | Subcortical shape/deformation | NA | FDR | right hemisphere only, for the accumbens, caudate, and putamen. |
| Burgaleta | 10.1016/j.neuroimage.2013.09.038. | 2014 | IQ change | 188 | Healthy adolescents 6-20 | Sex, scanner, time to repeat IQ | Cortical thickness, area | 20mm (thick) 40mm (area) | Sufstat 5,000 permutations (Nichols and Holmes, 2002) | 3 SA regions |
| Walhovd | 10.1016/j.neuroimage.2006.01.011 | 2006 | Memory recall (5 mins, 30 mins, 83 days) | 71 | Healthy adults 40+ | gender, age, IQ, and intracranial volume, (hippo volume) | Cortical thickness | 12.6mm | Uncorrected | Un-interpretable |
| Voineskos | 10.1002/hbm.22825 | 2015 | Cognition (verbal episodic memory, visuospatial episodic memory, and working memory) | 137 | Healthy adults 18-86 | Age, sex, education, APOE e4 status | Hippocampus shape (normalised fro TBV) | NA | 10% FDR (and 5%) (Genovese et al.,) | No significant associations at FDR <5% |
| Winjen | 10.1007/s00330-019-06437-9 | 2019 | EDSS and cognition domains | 34 | relapsing-remitting multiple sclerosis | Age, ICV | T1, T2, T2*, PD in grey matter masks | 10mm | Monte carlo, p<0.05 | T2 associations (no multiple correction for number of phenotypes studied) |

| | | | | | | | | | | |
|---|---|---|---|---|---|---|---|---|---|---|
| Bobholz | 10.1007/s11682-018-0005-z | 2019 | Cognition domains focus: psychomotor speed (digit symobol) | 135 | 81 idiopathic epilepsies<br>54 healthy controls | age, gender, and IQ, (epilepsy) | CV, CT, SA, and LGI (local gyrification index) | 15mm | Use of Qdec's Monte Carlo simulation allowed for corrections of multiple comparisons, with the cluster forming threshold set to p < 0.05 | LGI associations: left postcentral gyrus, left lateral occipital gyrus, and right caudal middle frontal gyrus |
| Brathen | 10.1002/hbm.24287 | 2018 | Episodic memory plasticity (improvment) | 126 | HC, did a 10weeks memory course<br>2 separate age groups | Age, sex, ICV | Cortical volume (FreeSurfer), fALFF | 15mm | The significance of this relationship was assessed within the FreeSurferframework (mri_glmfit), using cluster-based inference to account for multiple comparisons. To verify the reliability of the findings, several cluster-forming thresholds were tested, ranging from p < .05 to p < .001 (all tests were two-sided). | No significant relationships were observed between memory improvement and surface-level/vertex-wise cortical volume or cortical fALFF.<br>Similarly, no relationships were found at the MNI voxel-level when investigating noncortical fALFF. |
